# Comprehensive profiling of liver x receptor splicing in triple negative breast cancer reveals existence of novel splice variants that are prognostic for survival

**DOI:** 10.1101/2021.06.09.445161

**Authors:** Priscilia Lianto, J. Bernadette Moore, Thomas A. Hughes, James L. Thorne

## Abstract

The liver x receptors (LXR) alpha and beta are ligand-responsive transcription factors that link homeostatic control of lipid metabolism with cancer pathophysiology and prognosis. LXR activity is elevated in triple negative breast cancer relative to other breast cancer subtypes, driving gene signatures associated with drug resistance and metastasis. The loci encoding LXRα and LXRβ produce multiple alternatively spliced proteins, but the true range of variants and their relevance to cancer remain poorly defined. Seven splice variants of LXRα or LXRβ were detected. Three have not been recorded previously and five were prognostic. High expression of full length LXRα was associated with shorter disease-free survival but splice variants harbouring truncations of the ligand binding domain were prognostic for improved survival. All LXRβ variants were associated with longer disease-free survival. Mechanistically, while full length LXRα positively correlated with target gene expression in primary samples, LXRβ was inversely correlated. We conclude that canonical LXRα function is an oncogenic driver of triple negative tumour pathophysiology that can be countered by high expression of truncated splice variants and/or full length LXRβ.

**Highlights:**

- Expression of full length LXRα is associated with shorter disease-free survival of triple negative breast cancer patients
- A systematic evaluation of cell lines and primary tumour samples indicates LXR splicing is extensive in breast cancer
- Confirmation of three new LXR splice variants at transcript and protein level
- Expression of full length LXRβ or LXRα splice variants that harbour truncated ligand binding domains are associated with better prognosis
- Expression of LXR target genes is positively correlated with LXRα in relapsed patients and inversely correlated with LXRβ in survivors.

## 1. Introduction

Breast cancer (BCa) is the most commonly diagnosed cancer in women in the UK and is the cause of ~600,000 cancer death worldwide [1]. The triple-negative BCa (TNBC) subtype has higher rates of recurrence in the first three years after prognosis and increased mortality rates [2]. TNBC is defined by low oestrogen (ER), progesterone (PR), and Her2 receptor expression. Due to this lack of expression of therapeutic molecular targets primary disease can only be systemically treated with cytotoxic chemotherapy and TNBC remains a cancer of unmet clinical need; novel targeted therapies are urgently needed.

Nutritional and epidemiological studies indicate that cholesterol metabolism may play a role in the aetiology and severity of TNBC [3–6]. The liver x receptors (LXRα/NR1H3; LXRβ/NR1H2) are homeostatic regulators of cholesterol metabolism and as such have been suggested as potential therapeutic targets. LXRα activation by hydroxylated cholesterol (oxysterols) has been linked to poor survival by driving multi-drug resistance in TNBC patients [7], and increased metastasis in mouse models [8, 9]. On the other hand, LXRβ activation by histamine conjugated oxysterols such as dendrogenin A can induce lethal autophagy in breast cancer [10]. Intriguingly, ER-negative and ER-positive BCa subtypes respond differently to LXR ligands [11] even though oxysterol concentrations are similar between BCa subtypes [12]. Collectively, these data imply that the differential control over LXR’s transcriptional regulation between subtypes is not simply at the level of ligand type or concentration.

LXRα and LXRβ are each thought to be translated from their own single main transcript variants, producing 447 and 460 amino acid proteins respectively [13, 14].

These ‘full-length’ isoforms harbour domains that provide distinct functions, including activation function 1 (AF1), hinge region (H), DNA binding domain (DBD) and ligand binding domain (LBD) [13], as is typical for many ligand dependent nuclear receptors.

Aside from the canonical full-length LXRα1, four additional LXRα splice variants have been reported in human cell lines or tissue samples to date [15, 16], and a single report found one alternative splice variant to the major LXRβ1 isoform [17]. These studies found alternative splicing of LXR disrupts the integrity of the DNA and ligand binding domains, rendering the proteins with reduced or even absent response to ligand [13, 14]. In the case of LXRβ, a single known splice variant functions as an RNA co-activator [17]. Genomic and proteomic databases, such as NCBI, TCGA Splicing Variants database (TSVdb), ENSEMBL, and UNIPROT indicate that both LXRα and LXRβ are far more extensively spliced than has been reported. An analysis of splicing in the nuclear receptor superfamily predicted LXRα has 62 different transcript variants [18], making it the most extensively spliced nuclear receptor.

However, experimental evidence for the existence of such an array of isoforms is lacking. Given the strong links between cholesterol metabolism, for which the LXRs play vital regulatory roles, and TNBC aetiology, the aim of this study was to describe the LXR splice repertoire and clinical significance in human triple negative breast cancer.

## 2. Materials and Methods

### 2.1 Systematic transcript variant analysis in public databases

The NR1H3/LXRα and NR1H2/LXRβ spliced variant sequences were assessed in NCBI and ENSEMBL databases. Sequence annotation records of LXRα and LXRβ spliced variant read from databases were stored in a feature library created with apE plasmid editor. This feature library can then be used to scan any nucleotide and/or amino acid sequence that are complementary to all alternative forms of the LXRα and/or LXRβ sequences recorded in this library. The number of nucleotides in each exon and intron were counted in ApE plasmid editor to determine the proportions for drawing squares and lines represented exons and introns. The schematic diagrams of LXR variant structures were drawn using AutoCAD (Autodesk, US) software are shown in **Fig1**, **SF1**, and naming convention used in this study is described in Supplementary Information Section 1: Isoform naming rationale. Variants are listed in **TableS1 (LXRα)** and **TableS2 (LXRβ).**

**Figure 1.**
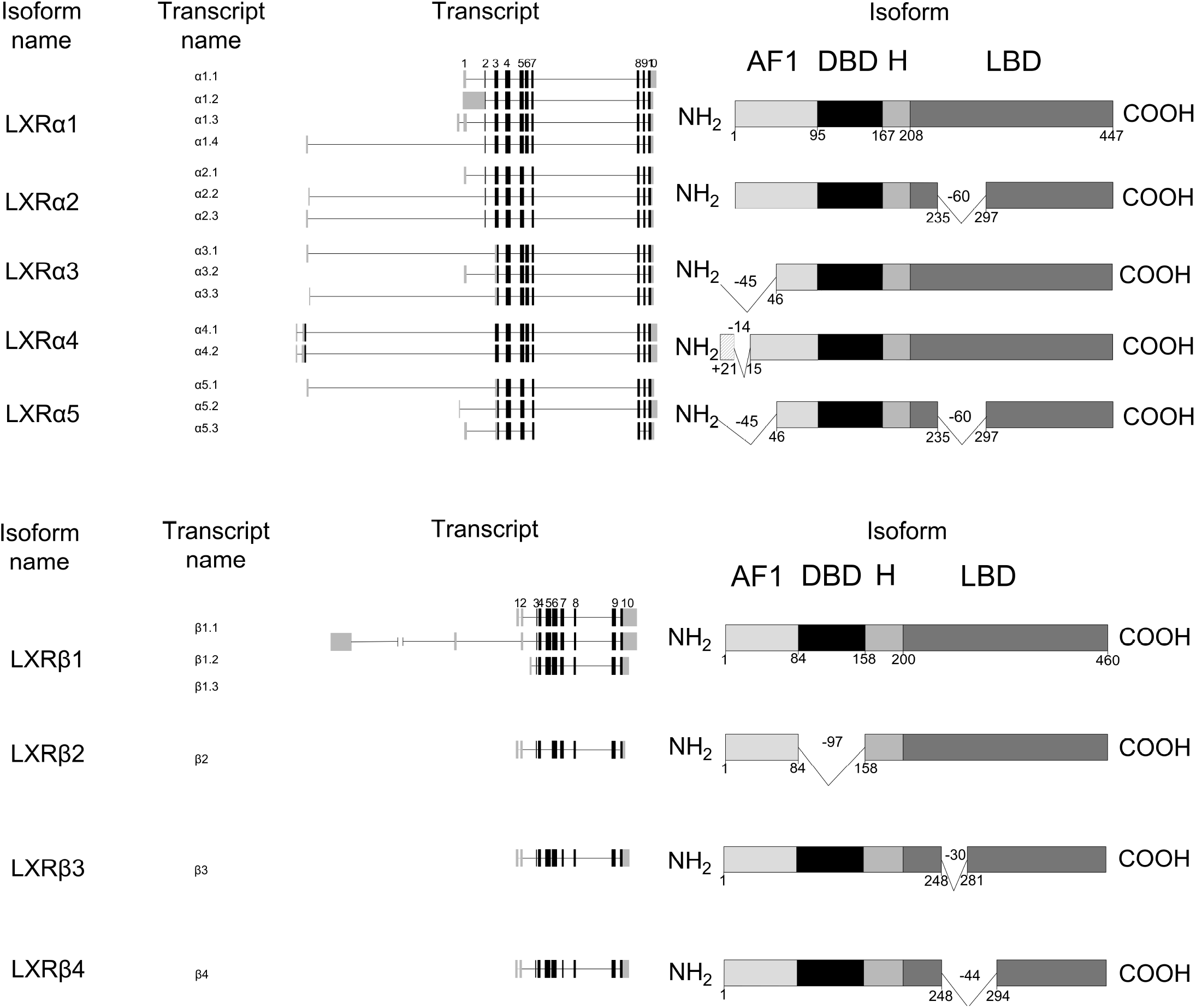
Schematic diagram of (A) LXRα and (B) LXRβ splice variants detected at RNA and protein level in this study. AF1=Activation Function 1; DBD=DNA Binding Domain; H=hinge; LBD=Ligand Binding Domain. The numbers right below the isoform domain boxes are represented the position of amino acids.

### 2.2 The Cancer Genome Atlas (TCGA) Splicing Database Analyses

TCGA splicing variants database (TSVdb) was searched and LXR splicing expression data in BCa tumour tissue, reported as normalised RNA-Seq by Expectation Maximisation (RSEM) values, were downloaded using the TSVdb webtool (**http://www.tsvdb.com**) [19]. Unreported and/or missing clinical status of the deposited TCGA BCa tumour samples were replenished by assessing patient tumour sample information from cBioportal (**http://cBioportal.org**) [20].

### 2.3 Human samples: ethical approval, collection, and processing

Thirty-eight frozen tumour samples were obtained with ethical approval from the Leeds Breast Research Tissue Bank (15/HY/0025 and LBTB_TAC_1/17). The clinicopathological features and selection criteria of these cohorts have been described previously [7, 11, 12, 21] (**Table-S3).** The Leeds TNBC tumour cohorts were grouped based on their disease-free survival (DFS) status. TNBC patients who did not have disease relapse during follow-up (median follow-up time was 96 months) were categorised to ‘No Event’ group. Meanwhile, TNBC patients who had relapsed and/or died due to their disease were grouped to an ‘Event’ group (median follow-up time was 20 months).

### 2.4 Cell culture

The HEPG2, MCF7, BT474, MDA.MB.231, MDA.MB.468, MDA.MB.453, and MDA.MB.157 cell lines were originally obtained from ATCC (Manassas, USA). Cell lines were routinely cultured in Dulbecco’s Modified Eagle Medium (DMEM; Thermo Fisher, UK, Cat. 31966047), supplemented with 10% Fetal Bovine Serum (FBS; Thermo Fisher, UK, Cat. 11560636) and incubated in 37°C, 5% CO2. Cells were periodically checked and confirmed to be mycoplasma free.

### 2.5 siRNA transfection

siRNA transfection was performed as previously reported [22]. Briefly, 1×10^5^ cells were plated in 6-well plates and incubated overnight. Lipofectamine RNAiMAX (Thermo Fisher, UK, Cat. 13778030), siLXRα (Origene, USA, Cat. SR322981) and siLXRβ (Origene, USA, Cat. SR305039) or the scrambled siRNA (Origene, USA, Cat. SR30004) were diluted in OptiMeM (Thermo Fisher, UK, Cat. 31985062), and added to the cells at a final concentration of 30 nM. The cells were incubated and after 20 h the media was removed and fresh DMEM added. Knockdown was confirmed at the protein level 48 h post-transfection.

### 2.6 mRNA extraction, cDNA synthesis, and qPCR

Total RNA was isolated from approximately 5×10^5^ cells using the ReliaPrepTM RNA Cell Miniprep System (Promega, UK, Cat. Z6012) following product guidelines, including on column digestion of gDNA. Cytoplasmic RNA was isolated from approximately 2×10^6^ cells using the Cytoplasmic & Nuclear RNA Purification Kit (Norgen, Belmont, CA, USA, Cat. 21000). The concentration of isolated RNA was determined spectrophotometrically, and RNA purity was evaluated by 260/230 nm and 260/280 nm ratios using a CLARIOstar plate reader (BMG LABTECH, Germany). cDNA was synthesised from 2 μg RNA using the GoScriptTM Reverse Transcription kit (Promega, UK, Cat. A5003). Primers were designed using NCBI BLAST primer design or, where transcript variants were not available in the NCBI database, primer 3 software was used [23]. Genomic DNA was digested, and cytoplasmic RNA was extracted to ensure intron retention was not confused with gDNA or incompletely spliced transcript. Amplicon size was restricted between 80 to 150 bp and amplicons were required to span an exon-exon boundary. Primer efficiency was measured from qPCR standard curves, and primers were redesigned if amplification efficiency did not fall within 90-110% and/or a single peak was not observed in melting temperature analysis. A description of primer design strategy is provided in Supplementary Information Section 2: Primer design strategy. Primer positions are shown in **SF2** and sequences provided in **Table-S4.**

### 2.7 Protein lysate extraction

Five mg fresh-frozen tumour biopsy samples were homogenised with a 1 mL Dounce tissue grinder (SLS, UK, Cat. HOM3580). Cell pellets and/or tumour samples were lysed in RIPA buffer (10 mM Tris-HCl pH 8, 140 mM NaCl, 0.1% SDS, 1% Triton X-100, 0.1% sodium deoxycholate, 1 mM EDTA, 0.5 mM EGTA), with 1 mM PMSF (Thermo Fisher, UK, Cat. 78440) added fresh prior to use. The lysed pellets were incubated on ice for 5 min and centrifuged at 11,500xg at 4°C for 10 min. The protein lysates concentrations were determined using a BCA kit (Thermo Fisher, UK, Cat. 23227) and lysates were kept in −80°C until further analyses. Each tumour sample was split in two and duplicate extractions were made. For each protein quantified (see 2.9) both duplicates were run and the average of each was used for analysis. Concordance of duplicates was high and is shown in **SF3**.

### 2.8 Cytoplasmic and Nuclear Protein Extraction

For the cytoplasmic and nuclear protein extractions, the REAP (Rapid Efficient And Practical) protocol [24] was followed with a slight modification. In brief, 1×10^7^ cells were suspended in 1 mL ice cold PBS with 1 mM PMSF. 200 μL of cell lysate was taken, put into a new chilled Eppendorf tube and centrifuged at 1,500 rpm for 3 min at 4°C. Supernatant was removed. The cell pellet (from 200 μL cell lysate) was resuspended in RIPA buffer with 1 mM PMSF (Thermo Fisher, UK, Cat. 78440) added fresh prior to use and labelled as a “whole cell fraction”. The remaining 800 μL cell lysate was centrifuged at 11,500xg at 4°C for 10 s. The supernatant was removed. The cell pellet (from 800 μL cell lysate) was resuspended in 200 μL ice-cold PBS+0.1 NP40 with 1 mM PMSF and centrifuged at 11,500xg at 4°C for 10 s. The supernatant was taken into a new chilled Eppendorf tube and labelled as a “cytoplasmic fraction”. The cell pellet was then resuspended in 200 μL ice-cold PBS+0.1 NP40 with 1 mM PMSF and centrifuged at 11,500xg at 4°C for 10 s. The supernatant was removed and cell pellet resuspended in 50 uL RIPA buffer with 1 mM PMSF. Cell lysate was sonicated with a water-bath sonicator (Diagenode Bioruptor Pico, Belgium, Germany) with cycles of 10 s on and 10 s off for five cycles, followed by centrifugation at 11,500 g at 4°C for 10 min. The supernatant was taken, put into a new chilled Eppendorf tube, and labelled as a “nuclear fraction”. The protein lysates concentrations were determined using a BCA kit and lysates were kept in −80°C until further analyses.

### 2.9 Immunoblotting

Forty-five micrograms of protein lysate combined with NUPAGE LDS sample loading buffer (Thermo Fisher, UK, Cat. NP0007) and DTT reducing agent (Thermo Fisher, UK, Cat. NP0004) was heated at 70°C for 10 min. For LXR variant expression, the protein lysate was loaded onto a 10% SDS polyacrylamide gel, electrophoresed at constant 80 V for 150 min, and transferred onto a PVDF membrane (Merck, UK, Cat. IPFL00010). The membrane was then blocked with TBS Odyssey Blocking Buffer (LI-COR Biosciences, UK, Cat. 92750000) for 1 h. Proteins were probed with anti-LXRα (R&D Systems, USA, Cat. PP-PPZ0412-00, dilution 1/1000), anti-LXRβ (Active Motif, Germany, Cat. 61177, dilution 1/1000), and anti-HPRT (Santa Cruz Biotechnology, Cat. sc-376938, dilution 1/100) overnight at 4°C. The membrane was then blocked and probed with LICOR secondary antibodies (IRDye 800CW goat anti-mouse Cat. 926-68170, IRDye 680RD goat anti-rabbit Cat. 926-68071; dilution 1/15,000, LI-COR Biosciences, UK) for 1 h and signal was visualised using the Odyssey system (LI-COR Biosciences, UK). Densitometry was performed using Image Studio™ Lite (LI-COR Biosciences, UK) software.

### 2.10 Immunoprecipitation

One mg protein sample was precleared with Dynabeads Protein A (Thermo Fisher, UK, Cat.100002D) and 1 μg isotype IgG2a (Cell Signalling, USA, Cat. 61656S). Immunoprecipitation was performed by incubating 40 μL of Dynabeads Protein A coupled to 2 μg of anti-IgG2a or 2μg of anti-LXRα using bis(sulfosuccinimidyl)suberate (Thermo Fisher, UK; Cat. A39266) with 1 mg protein sample overnight at 4°C. Dynabeads were washed 3 times with 10 mM Tris-HCl, 50 mM KCl (pH 7.5) and sample eluted in 20 μL of NUPAGE LDS sample loading buffer containing 100 mM DTT, then heated at 70°C for 10 min 10 at 70°C. The supernatant was transferred to a new Eppendorf tube after being separated from Dynabeads using a magnetic separator (Promega, UK, Cat. CD4002).

### 2.11 In-silico peptide mass prediction

The amino acid sequences of LXR variants downloaded from NCBI, ENSEMBL, and/or UNIPROT databases were subjected to the PeptideMass tool (http://www.expasy.org/tools/peptide-mass.html; Swiss Institute of Bioinformatics, Switzerland) [25] to theoretically trypsin digest the protein sequence in silico. A set of peptides from each LXR variant were then subjected to Diagram Venn (http://bioinformatics.psb.ugent.be/webtools/Venn; Bioinformatics & Evolutionary Genomics, Belgium) in order to find the unique peptides of each LXR variant.

### 2.12 S-Trap column coupled Mass Spectrometry (MS)

Protein samples were processed using the S-TRAP Micro column (PROTIFI, NY, USA) following the manufacturer’s instructions. Proteins were fully solubilised by adding 20 μL of 10% SDS solution to 20 μL sample in RIPA buffer. Reduction and alkylation were then performed. DTT was added to a final concentration of 20 mM before heating to 56 °C for 15 min with shaking. The sample was left to cool for 5 min then iodoacetamide was added to a final concentration of 40 mM, before heating to 20 °C for 15 min with shaking in the dark. Phosphoric acid was then added to a final concentration of 1.2%, to ensure destruction of all enzymatic activity and maximise sensitivity to proteolysis. Samples were then diluted with S-Trap binding buffer (100 mM TEAB pH 7.1 in methanol), and 1ug of trypsin, reconstituted in 50mM triethylamonium bicarbonate (TEAB), was added before the sample was quickly loaded onto the S-trap column. Proteins were captured within the submicron pores of the three-dimensional trap. Proteins captured within the trap present exceptionally high surface area allowing them to be washed free of contaminants. The S-trap was washed by adding 150 μL binding buffer before being spun at 4000xg for 30 s. 30 μL of 0.02 μg/μL Trypsin (Promega, WI, USA) was then added to the top of the S-trap. S-traps were loosely capped and placed in a 1.5mL Eppendorf and heated to 46 °C for 15 min with no shaking. Digested peptides were eluted by first spinning the S-trap at 4,000 g for 1 min. Further elution was performed in 40 μL 50mM TEAB, 40 μL 0.2% formic acid, and 30 μL 50% acetonitrile with 0.2% formic acid prior to centrifugation. Eluates were combined then dried down prior to resuspension in 0.2% formic acid.

### 2.13 Statistical analyses

All statistical analyses were performed using GraphPad Prism v8. The expression of LXR splice variants in TCGA tumours compared with the adjacent normal tissues was assessed using a Mann-Whitney two-tailed U test. Differential LXR splicing expression levels in both TCGA BCa and Leeds TNBC tumour samples were established using multiple t-tests with Holm-Sidak for multiple correction. The knockdown siRNA experiment was analysed using a two-tailed one-way ANOVA. The relationship between protein isoform variants and their mRNA transcripts were assessed using Spearman’s correlation and linear regression. ROC curves were used to establish the expression cut offs for high and low expression levels of each LXR variant. Tumour expression of each LXR variant was then assessed alongside patient survival to assess whether expression is predictive of survival in Kaplan Meier graphs. Patient survival was analysed using log-rank test.

## 3. Results

### 3.1 Database analysis of LXR transcript splice variance in breast cancer

To evaluate the variety of LXR splice variants, the NCBI and ENSEMBL databases were mined for mRNA transcript variants and UNIPROT database for protein variants. In total, this analysis indicated there were 64 LXRα transcript variants, of which 48 could code for 26 LXRα protein variants. We found 11 LXRβ transcript variants that could code for 9 LXRβ proteins (**SF1**). These included all nine variants (α1-α5, β1-β4) later observed in breast cancer cells and/or breast tumour tissue (summarised in **Fig1**).

To investigate the expression of LXR splice variants in BCa, we first examined RNA-seq data from 1,103 BCa tumours in the TCGA Pan-Cancer Atlas utilising the TSVdb web-interface [19]. These data showed that out of six LXRα and three LXRβ splice variants with RSEM reads [26], the α1.1 (median RSEM value=231.52) and β1.1 (median RSEM value =1158.53) variants were most highly expressed in primary BCa samples (**Fig2A**). The α3.1 (median RSEM value=115.42), α1.2 (median RSEM value=31.88) and β2 (median RSEM value=21.69) transcripts were expressed at low levels. No other transcript variants were detected in this database (**Fig2A**).

**Figure 2.**
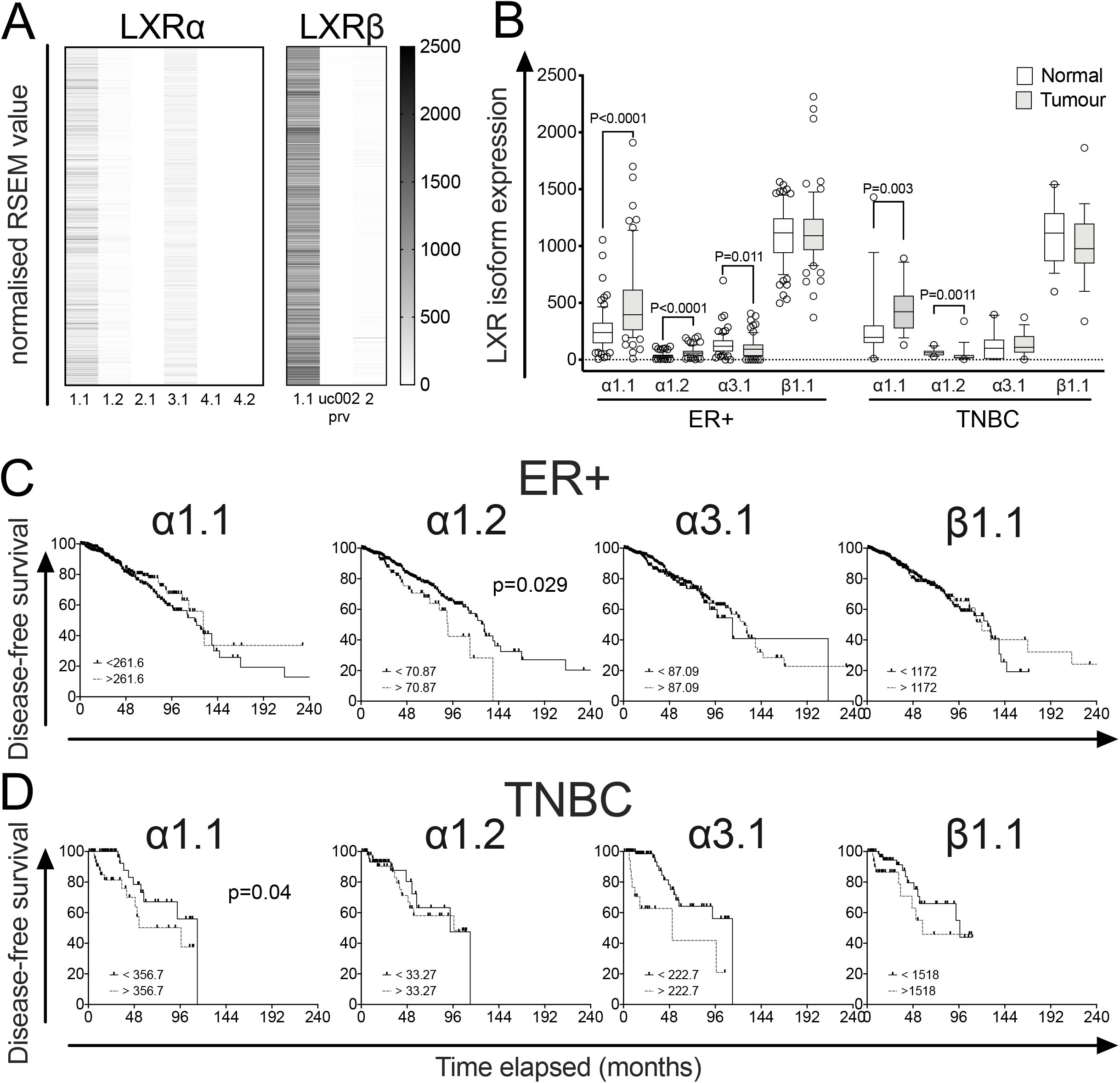
LXR Splicing events in BCa validated by TCGA Splicing Variants database (TSVdb). (A) Heatmap visualization of normalized RSEM (RNA-Seq by Expectation Maximization) value expression of LXRα and LXRβ in 1,103 TCGA BCa samples. Note: Transcript uc002prv was only reported in TSVdb and is not recorded in NCBI, ENSEMBL, or UNIPROT databases. (B) Comparison of LXR transcript variant expression, for α1.1, α1.2, α3.1, and β1.1, in matched tumour and normal tissues grouped by BCa subtype, ER+ (n=78) and TNBC (n=18). Statistical analysis by Man-Whitney two tailed U tests. P≤0.05 was considered significant. (C, D) Kaplan-Meier survival curves plotting disease free survival of TCGA BCa patients. The TCGA BCa samples (downloaded from TSvdb TCGA splicing database) were grouped by BCa subtype, (C) ER+ (n=803) and (D) TNBC (n=101). Survival curves of each subtype were separated into two groups, no event (ER+=675, TNBC=79) and event (ER+=128; TNBC=22) based on their overall survival data reported in cBioportal. P-value is based on the log-rank test and data groups were considered significantly different if p<0.05.

We further examined expression of the four variants detected at highest levels in matched tumour and normal samples from ER+ and TNBC subtypes. Both α1.1 and α1.2 were expressed at significantly higher levels in ER+ and TNBC tumour tissue compared to adjacent normal tissue (P<0.01 for all; **Fig2B**). α3.1 expression was lower in ER+ tumour tissue than adjacent normal (P=0.011; **Fig2B**) but there was no difference in expression of this isoform between tumour and normal tissue in TNBC disease. There was no difference in β1.1 expression in either ER+ or TNBC tumours relative to adjacent normal (P>0.05; **Fig2B**). There was no difference in expression of any individual LXR variants between ER+ and TNBC tumours (**SF4**).

We then examined if transcript variant expression identified in TSVdb was associated with disease-free survival (DFS) in ER+ (**Fig2C**) or TNBC patients (**Fig2D**). High α1.2 expression was associated with shorter DFS in ER+ patients (p=0.029), while in TNBC patients, high α1.1 expression was associated with shorter DFS (p=0.04). Neither α3.1 or LXRβ expression was associated with DFS. These data suggest that elevated LXRα1 (a1) may be linked to poor prognosis in breast cancer patients.

### 3.2 LXR expression in breast cancer cell lines

Having established evidence for LXR splice variant expression in the TSVdb dataset, we next examined transcript expression in a panel of BCa cell lines. HEPG2 cells express relatively high levels of both LXRα and LXRβ [27] so were included as a positive control. Total LXRα and LXRβ RNA expression was determined using SYBR green primers that target the exon 9-10 junction of LXRα and exon 8-9 junction for LXRβ. These exons are spliced together in every previously reported variants and in all variants described in this study **(Table-S1+S2; SF1-A-B)** and were used to estimate the total pool of LXR transcripts and for normalisation of variant expression.

The claudin high TNBC cell line MDA.MB.468 had highest expression of LXRα of all BCa cell lines (P<0.05). MDA.MB.468, MDA.MB.453 and BT474 had highest LXRβ (**Fig3A-B**). In all cell lines analysed, the expression of LXRα mRNA was significantly lower than LXRβ (all p<0.05; **SF5**), recapitulating the observations from the TSVdb (**Fig2A**). Across the cell lines, the transcripts measured accounted for between 91-100% of LXRα (exon 9-10) and 94-102% of LXRβ (exon 8-9) exon-exon boundaries (**Fig3C-D**), indicating other variants were expressed at very low levels or did not contain the exon 9-10/7-8 boundaries. Across all cell lines we detected five mRNA species (α1.1, α1.3, α2.3, α3.1, α5.3; **Fig3C**) predicted to code for four LXRα protein variants (α1, α2, α3, α5), and two coding for LXRβ (β1, β4; **Fig3D**). LXRα5 and LXRβ4 have not been reported in the literature previously.

**Figure 3.**
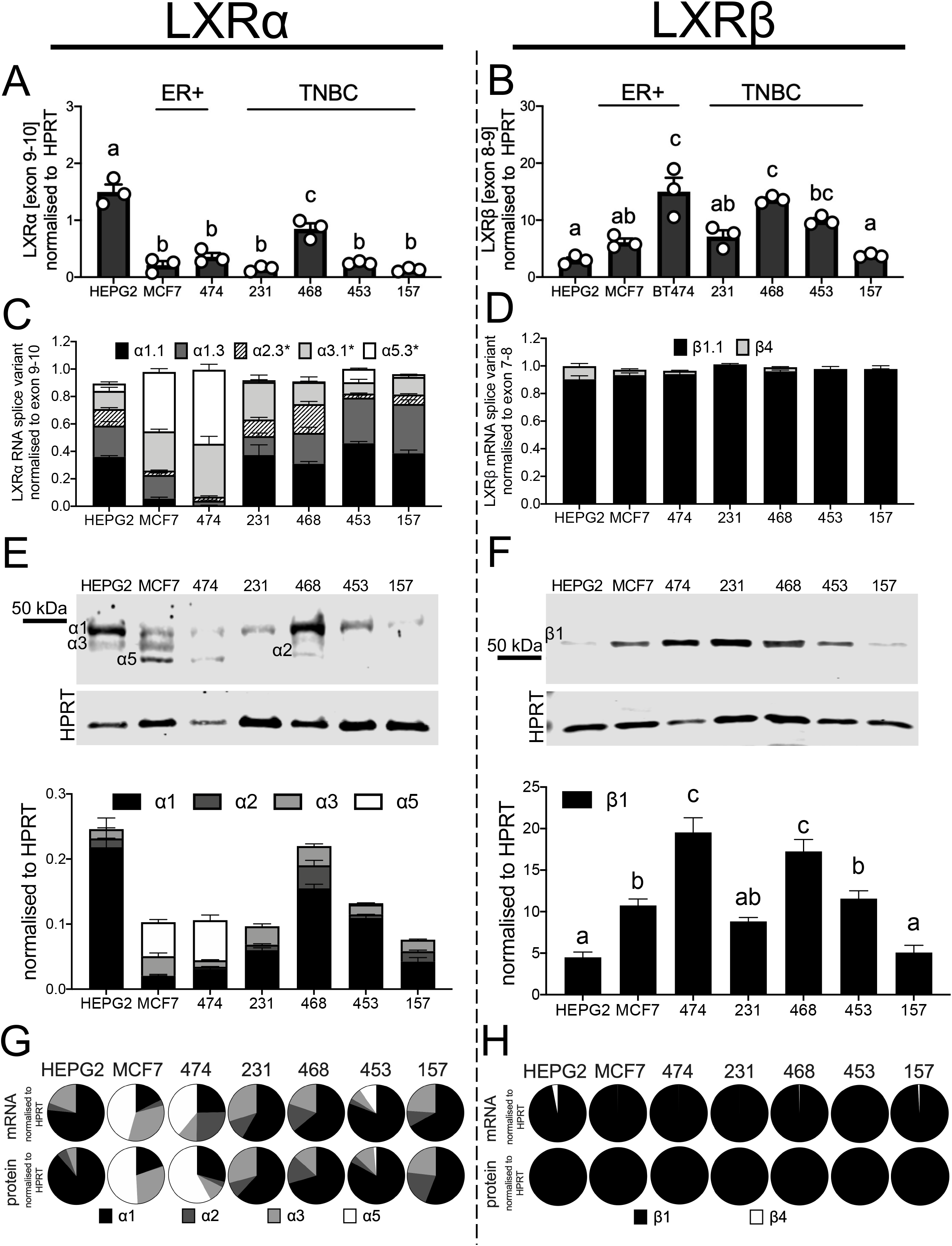
Diversity of LXR splice variants in breast cancer cell line panel. RNA analyses of total LXRα (A) and LXRβ (B) expression. The stacked bar graphs show LXRα (C) and LXRβ (D) transcript variants normalised to ubiquitously expressed exon-exon junctions. Asterix denote ambiguous amplicons. Representative blots and densitometry of LXRα (E) and LXRβ (F) protein variants observed in cell lines. Pie charts show contribution of each transcript variant and protein variant to the total amount of LXRα (G) and LXRβ (H). All data shown are mean of three independent replicates with SEM. Statistical significance was measured by two-tailed one-way ANOVA and significant differences are denoted with different letters if p≤0.05.

α1 transcripts were dominant (50-80% of all LXRα) in HEPG2 and TNBC cells (**Fig3C**: note black and dark grey sections corresponding to α1.1 and α1.3 respectively). In the ER+ cell lines α5 (40-50%) and α3 (30-40%) were the majority species (**Fig3C**: note light grey and white sections corresponding to α5.3 and α3.1 respectively). α2 comprised between 5-20% depending on the cell line (**Fig3C**: hatched sections). LXRβ1.1 was detected in all BCa cell lines, with a very small amount of β4 transcript found in HEPG2 and the ER+ cell lines only (**Fig3D**). We successfully detected over 90-100% of all the LXR variants (harbouring the exon 9-10 and 7-8 junctions) present in the cell lines. From these data, we concluded that full-length α1 was the dominantly expressed variant in TNBC cells, truncated α5 was the dominant LXRα isoform in ER positive cells, and β1 was the dominant LXRβ transcript across all cell types.

We next performed immunoblotting to establish the range of protein variants present. Representative blots and densitometry analysis show bands corresponding to predicted sizes of LXRα1 (≈50kDa) and LXRβ1 (≈55kDa) were robustly and reproducibly detected in all cell lines (**Fig3E** and **Fig3F** respectively). However, optimisation of immunoblotting experimental conditions, including antibody choice, indicated multiple additional bands were present when probing for LXRα (**Fig3E**). These bands closely corresponded to sizes of proteins that would be coded by the transcripts identified above, namely: α1 (50 kDa), α2 (44 kDa), α3 (46 kDa), and α5 (39 kDa). The cell line specific pattern of protein expression matched that of the transcript expression; a1 RNA and protein were highly expressed in HEPG2 and the TNBC cell lines while a5 was dominant in the ER+ cells (**Fig3G**). For LXRβ only a single isoform β1 was present at both RNA and protein level (**Fig3H**).

We concluded that in TNBC cell lines full-length LXRα1 is the most abundant isoform comprising 50-80% of LXRα protein (depending on cell line), with two a1 transcript variants accounting for 51-75% of LXRα transcript. LXRα3, which lacks part of the AF-1 domain and has diminished response to ligand [15] was the next most abundant producing 15-30% of the protein and 8-27% of the transcript. LXRα2, which is non-responsive to ligand [15], made up 8-20% of the total LXRα protein coded by two transcripts producing 7-12% of the LXRα coding RNA (see **Supplementary Information Section 1** for variant nomenclature). Only a small amount of LXRα5 was detected in TNBC (<10% in MDA.MB.453 cells and undetected in other TNBC lines). Interestingly, this ‘double-truncation’ variant that lacks both the AF-1 region of a3 and the LBD region of α2, was the most abundant isoform in ER+ cells at protein (35-70%) and transcript level (43-54%).

### 3.3 LXR splice variant expression and clinical significance in triple negative breast cancer

To evaluate LXR variant expression and significance in clinical samples, LXR protein expression was measured in a cohort of 38 fresh-frozen TNBC tumour samples (mean follow-up of three years), which has been reported on previously [11, 12] (Table-S3). Patients were dichotomised into two groups: ‘No Event’ patients defined as those who were alive and disease free at the time of last reporting or had died from unrelated reasons; ‘Event’ patients were those who had died from their disease and/or had disease recurrence. Representative blots are shown in **Fig4A**. Event patients had significantly higher expression than No Event patients of both full-length isoforms (α1: p<0.0001; α4: p=0.001; **Fig4B**). Together α1 and α4 comprised 70% of the total LXRα protein in the No Event group, but this rose to 93% in the Event group (**Fig4C**). No significant difference was observed in level of α2, α3, α5, β1, or β4 protein variants between groups (**Fig4B**).

**Figure 4.**
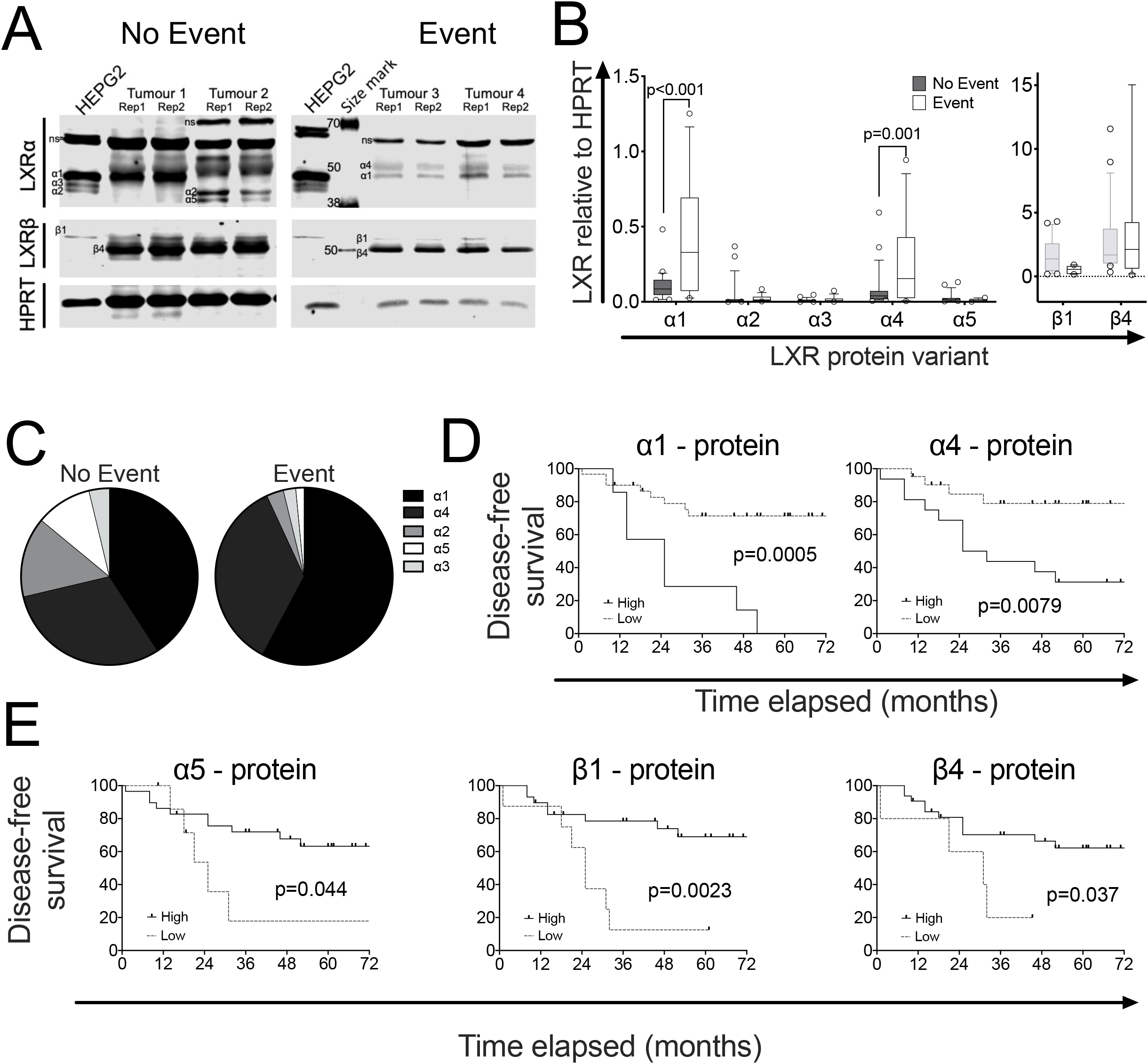
LXRα1 and LXRα4 are highly expressed in TNBC patients with poor survival. Representative blots showing the LXR splicing events in tumours derived from 38 TNBC patients, who were either free from disease (no event; n=23) or who had relapsed or died (event; n=15) after follow-up (A). Differential expression of LXRα and LXRβ protein variants in No Event and Event groups were plotted in box and whiskers charts (B). The line shows the median value, the box shows 10th to 90th percentile and whiskers show minimum to maximum values. Statistical analysis was established using multiple *t*-tests with Holm-Sidak for multiple correction, p≤0.05 was considered significant. Pie charts show contribution of each transcript variant and protein variant to the total amount of LXRα in No Event and Event patients (C). Kaplan-Meier survival curves plotting disease free survival of TNBC patients dichotomised based on protein expression of full length LXRα1 and LXRα4 transcripts (D), or truncated LXRα5 and LXRβ1 and LXRβ4 (E). All protein measured relative to HPRT. Data derived from the mean of two different slices of tumours. Significance determined by the Log-rank (Mantel-cox) test where p≤0.05 was considered significant.

When patients were dichotomised using ROC analysis, high protein expression of the full-length variants were, as expected, found associated with significantly shorter disease-free survival (α1: p=0.0005; α4: p=0.0079; **Fig4D**). Interestingly, in this more nuanced analysis, high expression of α5 (p=0.044), LXRβ1 (p=0.0023), and LXRβ4 (p=0.037) was associated with significantly longer DFS (**Fig4E**). These observations were replicated when dichotomising based on RNA expression (**SF6A**). Furthermore, a3.1, which codes for intact LBD but harbouring a deletion in the AF1 domain, was also associated with shorter DFS (p=0.022; **SF6A**). The a2.3 transcript, which codes for a protein with the same LBD deletion as a5 was, like a5, associated with longer DFS (p=0.031; **SF6A**). a2 and a3 protein however were not associated with DFS (**SF6B**). We concluded that full length LXRα isoforms may exacerbate disease severity, whereas those lacking the full LBD or LXRβ isoforms were associated with reduced disease severity.

### 3.4 Validation of isoform identity

#### 3.4.1 Bands representing LXR variants were reduced by targeted siRNA

To confirm if the protein bands were LXRα variants, we first performed siLXRα treatments in cell lines. Using previously validated [7] siRNA duplexes (Origene trisilencers) against LXRα and LXRβ we found that the protein bands predicted by size to be α1, α2, α3, a5, and β1 were all significantly reduced by targeted siRNA in all cell lines tested (all p<0.05; **SF7**). Note: a4 and β4 were not expressed in cell lines so could not be validated in this way. Furthermore, siLXRα did not reduce LXRβ expression and siLXRβ did not reduce expression of any LXRα isoform, and, as is characteristic of LXRs [15, 28], all protein variants were localised in the nucleus (**SF8**).

#### 3.4.2 Unique peptides representing a4 and β1 were identified by MS S-trap Mass Spectrophotometry

As was the case for cell line analyses, confirmation of the identity of the observed protein variants in tumour tissue was important, especially having identified the α4 variant which has not previously been reported in the literature. As siRNA was not possible on the tumour samples, we performed S-trap [29] coupled mass spectrometry (St-MS) in cell line and tumour lysates samples selected to represent as much of the diversity in isoforms as possible (**Table1**, **Table2**, **ST5**, **ST6**). This included two tumours in duplicate, ligand treated MDA.MB.468 cells (**SF9A**), and in siCON, siLXRα, siLXRβ (**SF9B**). We found 15 peptides uniquely corresponding to α4 BLAST sequence alignment indicated they were high-quality hits (100% identity), including the two first exons coding for the alternative AF1 domain in this isoform (**SF9C** – boxed amino acids) supporting our prior conclusion that α4 was expressed in some tumour samples. Unique peptides of β1 were also detectable in our tumour sample (**SF9D** – boxed amino acids). However, unique peptides corresponding to α1, α2, α3, α5, or β4 were not produced by our enzymatic cleavage and in silico analysis indicated these isoforms would not be distinguishable from each other (**Table3).** Our next approach was to perform immunoprecipitation. We found this successfully enriched each isoform (**SF9E**) but limited amounts of tumour meant bands could not be visualised for excision on coomassie gels. We concluded that α4 and β1 were present in tumour tissue.

**Table 1.**
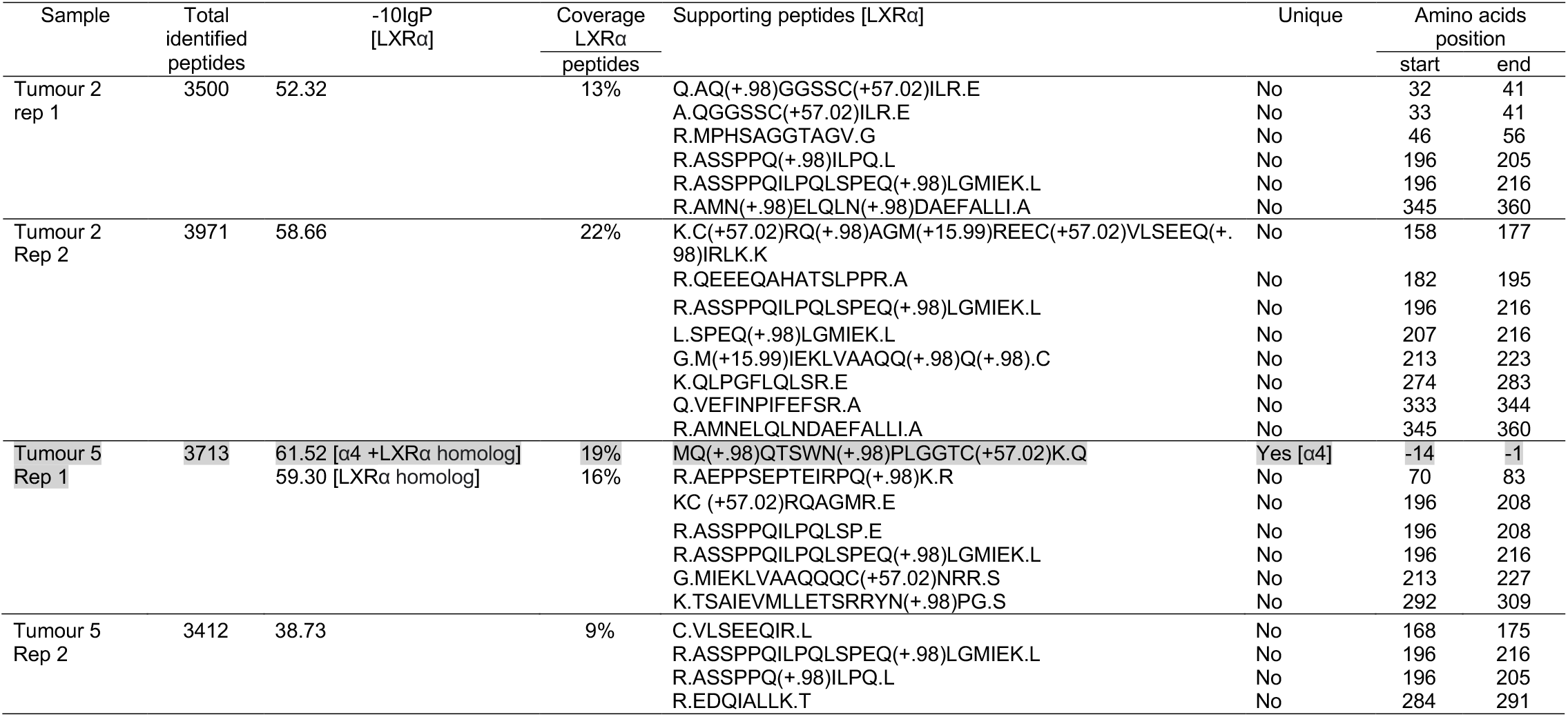
LXRα peptides detected by S-trap column coupled with Mass Spectrometry (MS). Amino acids position numbers based on LXRα1 structure. The grey highlight indicated unique peptides of LXR variant detected by MS.

**Table 2.**
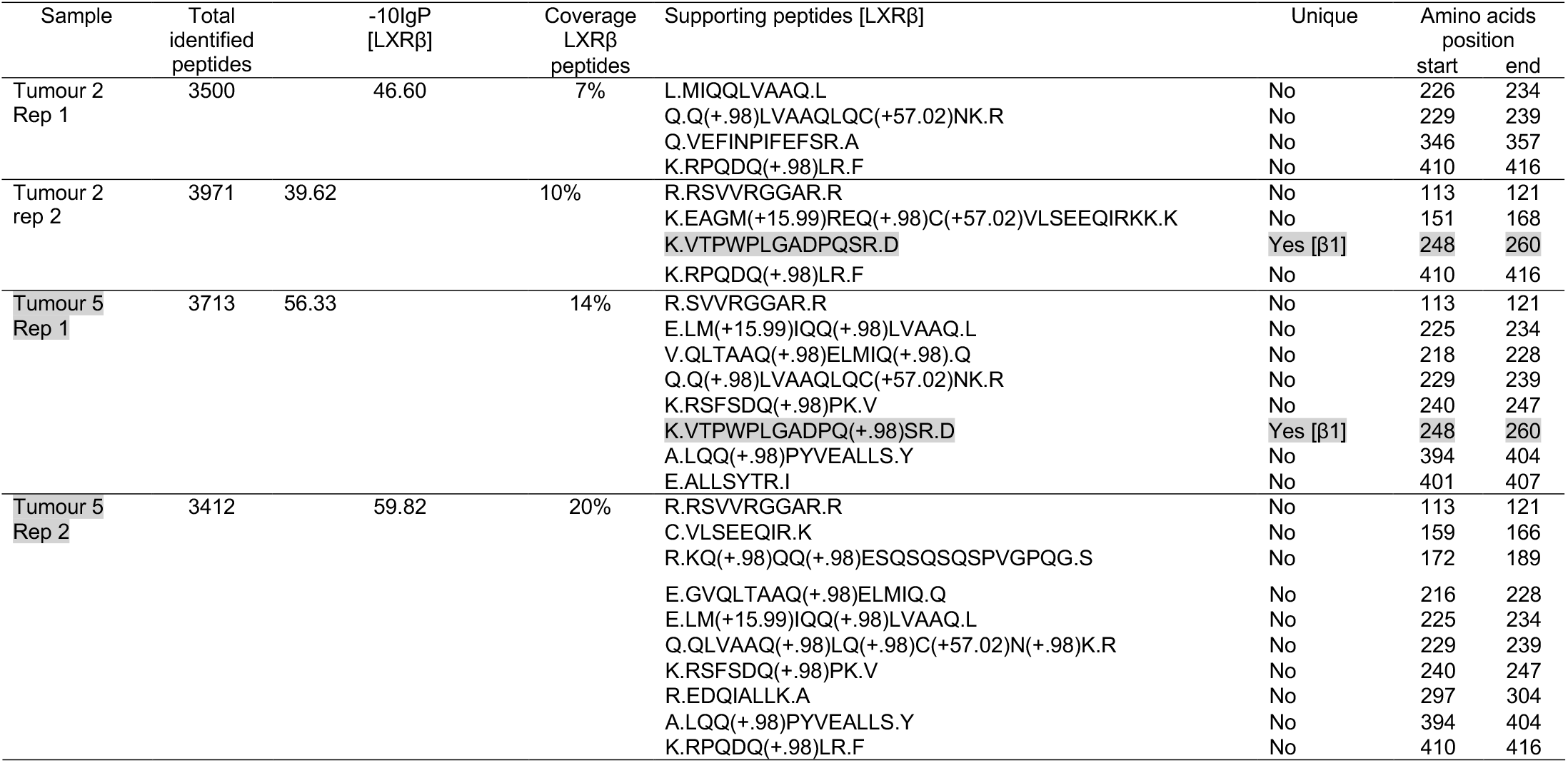
LXRβ peptides detected by S-trap column coupled with Mass Spectrometry (MS). Amino acids position numbers based on LXRβ1 structure. The grey highlight indicated unique peptides of LXR variant detected by MS.

| Sample | Total identified peptides | -10lgP [LXRβ] | Coverage LXRβ peptides | Supporting peptides [LXRβ] | Unique | Amino acids position | |
| --- | --- | --- | --- | --- | --- | --- | --- |
|  |  |  |  |  |  | start | end |
| Tumour 2<br>Rep 1 | 3500 | 46.60 | 7% | L.MIQQLVAAQ.L | No | 226 | 234 |
|  |  |  |  | Q.Q(+.98)LVAAQLQC(+57.02)NK.R | No | 229 | 239 |
|  |  |  |  | Q.VEFINPIFEFSR.A | No | 346 | 357 |
|  |  |  |  | K.RPQDQ(+.98)LR.F | No | 410 | 416 |
| Tumour 2<br>rep 2 | 3971 | 39.62 | 10% | R.RSVVRGGAR.R | No | 113 | 121 |
|  |  |  |  | K.EAGM(+15.99)REQ(+.98)C(+57.02)VLSEEQIRKK.K | No | 151 | 168 |
| | | | | K.VTPWPLGADPQSR.D | Yes [β1] | 248 | 260 |
|  |  |  |  | K.RPQDQ(+.98)LR.F | No | 410 | 416 |
| Tumour 5<br>Rep 1 | 3713 | 56.33 | 14% | R.SVVRGGAR.R | No | 113 | 121 |
|  |  |  |  | E.LM(+15.99)IQQ(+.98)LVAAQ.L | No | 225 | 234 |
|  |  |  |  | V.QLTAAQ(+.98)ELMIQ(+.98).Q | No | 218 | 228 |
|  |  |  |  | Q.Q(+.98)LVAAQLQC(+57.02)NK.R | No | 229 | 239 |
|  |  |  |  | K.RSFSDQ(+.98)PK.V | No | 240 | 247 |
| | | | | K.VTPWPLGADPQ(+.98)SR.D | Yes [β1] | 248 | 260 |
|  |  |  |  | A.LQQ(+.98)PYVEALLS.Y | No | 394 | 404 |
|  |  |  |  | E.ALLSYTR.I | No | 401 | 407 |
| Tumour 5<br>Rep 2 | 3412 | 59.82 | 20% | R.RSVVRGGAR.R | No | 113 | 121 |
|  |  |  |  | C.VLSEEQIR.K | No | 159 | 166 |
|  |  |  |  | R.KQ(+.98)QQ(+.98)ESQSQSQSPVGPQG.S | No | 172 | 189 |

|  |  |  |  |
| --- | --- | --- | --- |
| E.GVQLTAAQ(+.98)ELMIQ.Q | No | 216 | 228 |
| E.LM(+15.99)IQQ(+.98)LVAAQ.L | No | 225 | 234 |
| Q.QLVAAQ(+.98)LQ(+.98)C(+57.02)N(+.98)K.R | No | 229 | 239 |
| K.RSFSDQ(+.98)PK.V | No | 240 | 247 |
| R.EDQIALLK.A | No | 297 | 304 |
| A.LQQ(+.98)PYVEALLS.Y | No | 394 | 404 |
| K.RPQDQ(+.98)LR.F | No | 410 | 416 |

**Table 3.**
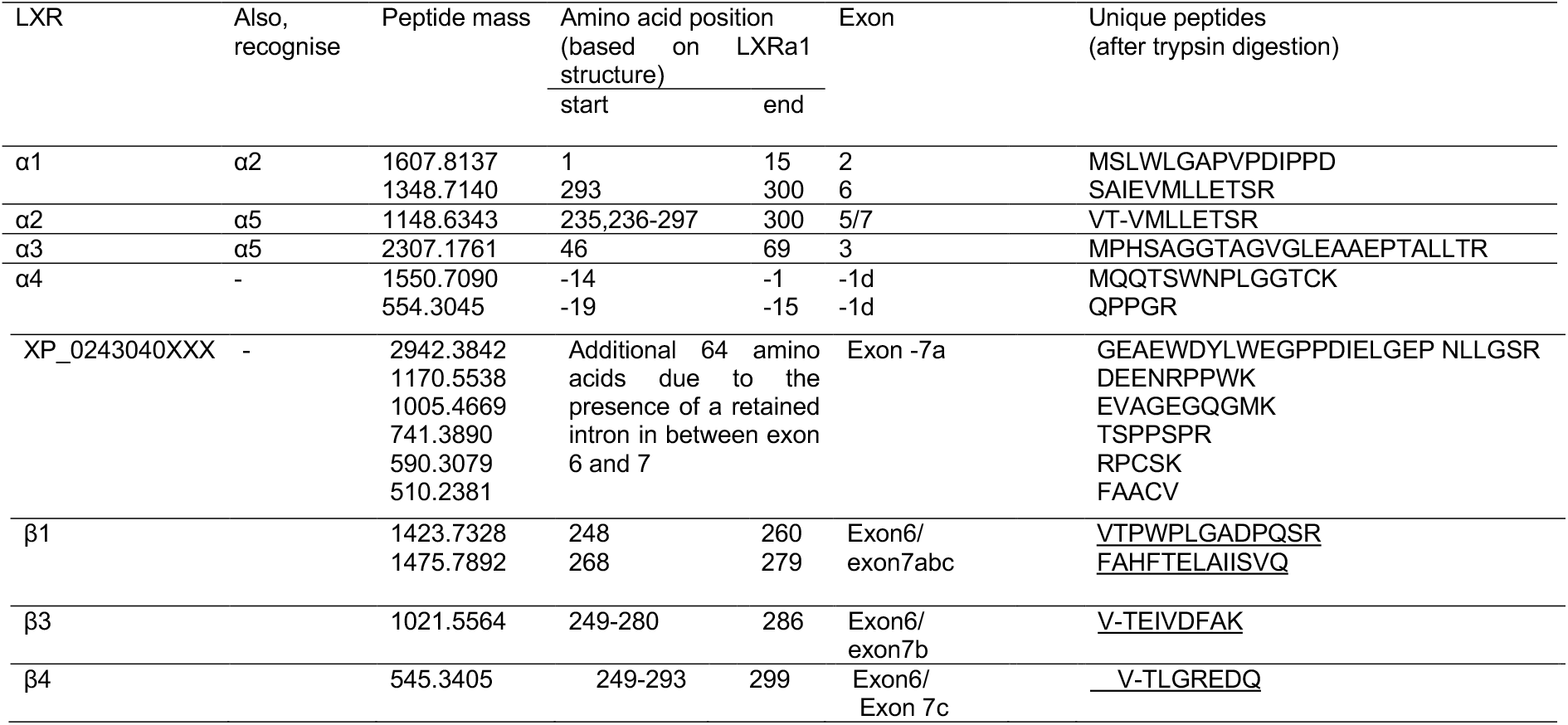
Unique peptides of LXR variants. Amino acids position number based on α1 and/or β1 amino acid numbering position. Amino acid numbering position with “-” indicates the additional amino acid(s) coming before α1 and/or β1’s amino acid position number 1. The numerical exons represent the first-discovered exons. The numerical exons followed with alphabets represent the later-discovered exons.

#### 3.4.3 Transcript-protein correlation analysis

Linear correlation analysis was performed in the seven cell lines comparing expression of each transcript variant against each protein band for which it could potentially code. There was a strong and significant positive correlation between transcript variants and the protein isoform for which they were predicted to code: a1 protein correlated with a1.1 (R^2^=0.97; p<0.0001; **SF10** top row) and a1.3 (R^2^=0.78; p=0.0084; **SF10** top row). β1 protein correlated with β1.1 transcript (R^2^=0.99; p<0.0001; **SF10** bottom row). During primer design it became clear that due to complexity arising from the large number of LXR coding transcripts, several transcripts were not distinguishable from each other due to sequence homology (**see Supplementary Information Section 1 and 2**, and denoted with asterixis in **Fig3C**). Specifically, indistinguishable transcripts were: a1.4 and a2.3; a3.1 and a5.1; a3.3 and α5.3. We tested all potential protein isoforms for correlation with the ambiguous primer pairs: a1.4/a2.3 correlated with a2 protein (p=0.0022; R^2^=0.97; **SF10** row 2) but not a1 protein (p>0.05); a3.1/a5.1 correlated with a3 protein (p=0.034; R^2^=0.63; **SF10** row 3) but not a5 protein (p>0.05); a3.3/a5.3 correlated with a5 protein (p=0.029; R^2^=0.65; **SF10** row 4) but not a3 protein (p>0.05). We also designed primers against the exon 5 and 7 junction to simultaneously recognise the a2 and a5 variants that lack exon 6 and a portion of the LBD. These a2/a5 primers detected RNA that correlated with a5 protein (p=0.0065; R^2^=0.8; **SF10** fourth row) but not with a2 (p>0.1; **SF10** second row).

Next, although cytoplasmic extraction was not possible in frozen tumour samples as they had lost cellular substructure in the freezing process, given the number of replicates was larger (n=38) we considered testing for correlations a reasonable approach (**SF11**). As for cell lines, both a1.1 and a1.3 correlated with a1 protein (p<0.05 for both); α2.3 correlated with α2 protein (p=0.036); α3.1 correlated with α3 protein (p=0.056); α4.1 correlated with α4 protein (p=0.0037; note, this variant was not detected in cell lines); α5.3 (p=0.039) and α2/α5 (p=0.054) correlated with α5 protein. For LXRβ, β1.1 transcript correlated with β1 protein (p=0.022). β3 and β4 are almost identical in size (46 and 47 kDa respectively) so are indistinguishable by immunoblotting. β4 transcript (p=0.031) but not β3 (p>0.05) corelated with the band corresponding to β3/β4 protein (**SF11** bottom row) confirming this as LXRβ4. In combination with the siRNA in the cell line studies above and MS experiments, these correlative observations indicate five LXRα variants and two LXRβ variants are expressed in TNBC.

### 3.5 LXR splice variants are differentially correlated with expression of target genes in Event and No Event patients

We previously reported that LXRα expression is positively correlated to target gene expression in ER-negative patients who had relapsed and/or died due to their disease (Event patients) but not those who survive disease free (No Event patients) [7]. Here, we tested the hypothesis that differential LXR splice variant expression between cancers may contribute to disease aetiology via their ability to control expression of gene targets. LXR splice variant expression was tested for correlations with established LXR target genes, ABCA1 and ABCB1 [7]. Expression of both full-length isoforms (α1 and α4) positively correlated with expression of both ABCA1 or ABCB1, but interestingly only in Event patients (p<0.05 for all; **Fig5**). Strikingly, expression of β1 was inversely correlated with both ABCA1 and ABCB1 but only in No Event patients (p<0.05; **Fig5**). Statistically significant correlations between α2, α3, or α5 with target genes were not observed (**SF12**). From these data we concluded that while full length LXRα is associated with activation of gene targets in patients who relapse or die, LXRβ is associated with inhibition of the same target genes in TNBC survivors.

**Figure 5.**
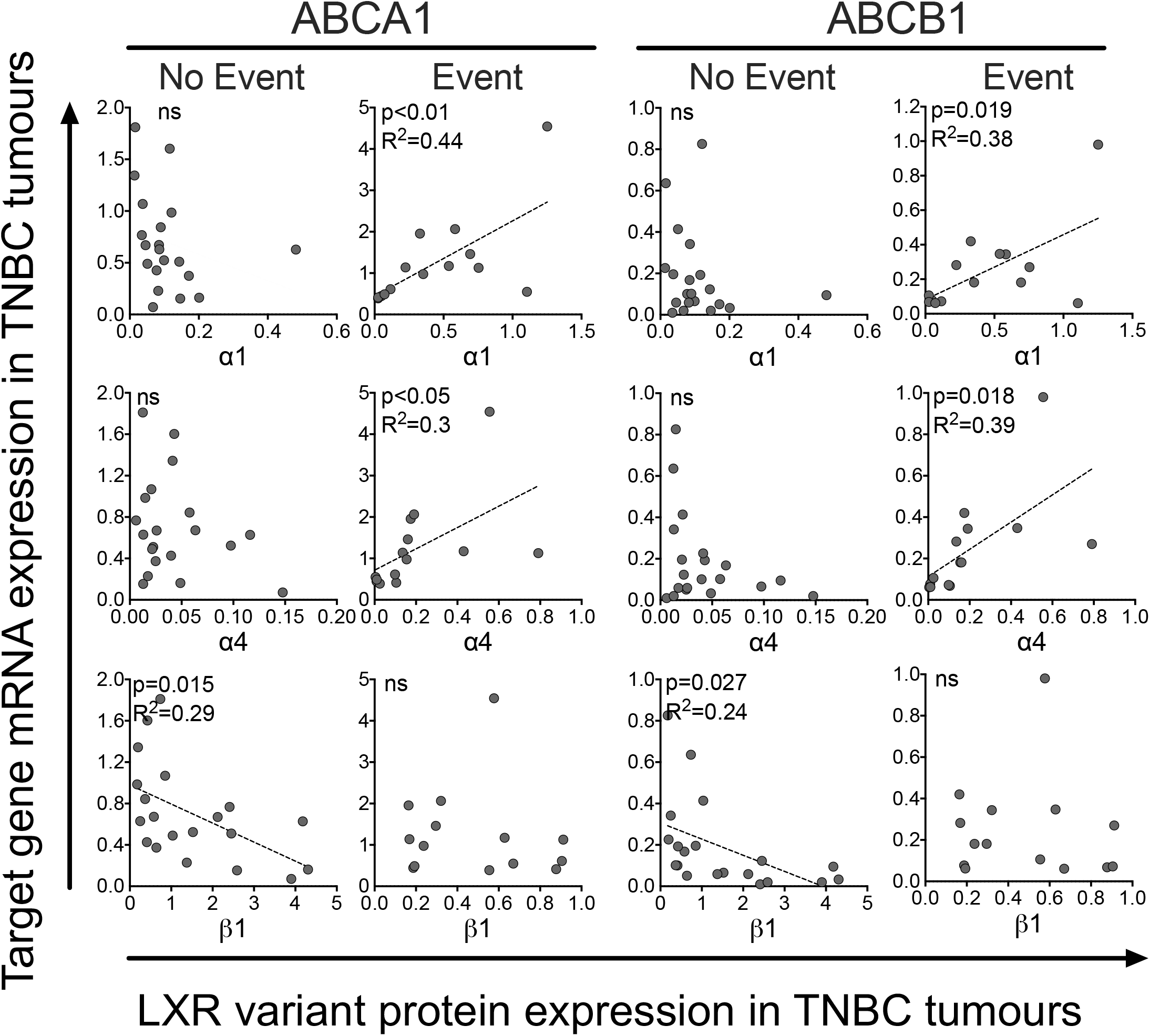
Protein expression of LXR variants is differentially associated with LXR target gene expression in TNBC tumours. TNBC patients (n=38) were divided into two groups, no event (n=23) and event (n=15) based on disease-free survival status. Linear regression was used to determine if the slope of the lines were significantly different from 0 (p≤0.05). Line of best fit are shown if the relationship was significant. Circles represent mean protein expression of two tumour slices for individual TNBC patients.

## Discussion

The objective of this study was to establish the repertoire, expression levels, and pathophysiological significance of LXR splice variants in breast cancer. In TNBC we found evidence that five LXRα and two LXRβ splice variants are expressed at RNA and protein levels. Three of these isoforms, α4, α5, and β1, are new to the literature and do not have validated UNIPROT records. Our data demonstrate that LXR splice variants have clinical significance in TNBC; full length LXRα (α1 and α4) is associated with shorter DFS, while LXRβ and LXRα variants with truncated ligand binding domains were associated with longer DFS.

We provide several lines of evidence that the LBD is required for LXRα’s oncogenic potential. Firstly, high expression of the full-length α1 was strongly associated with shorter DFS at both transcript and protein level. High expression of two other isoforms, α3 (transcript only) and α4 (transcript and protein) were also associated with shorter DFS. Although both have disruptions to their AF1 domains, they are otherwise homologous to full length α1, including, importantly, an uninterrupted LBD. α5 skips the same exons as α2 and α3 and has the same deletions; 60 amino acids are missing from the LBD, and 45 amino acids of the AF1 domain are missing, respectively. These truncations have been shown separately to diminish (a3) or completely abolish (a2) transcriptional activity [15]. Interestingly, high transcript expression of a2 was, like a5, linked to longer DFS. High a3 transcript on the other hand, like full length a1 and a4 was linked to shorter DFS. These observations suggest that splice variants with disrupted LBD are advantageous, but the protein can tolerate disruption or even partial deletion of the AF1 domain and still be associated with worse prognosis. β1 (P55055-1) and β4 (M0R2F9) are generated by the mutually exclusive inclusion or exclusion of exon 7, again leading to a change in the LBD. With LXRβ however, high expression of either β1 and β4 was linked to longer DFS. The cellular role of this altered LBD remains to be determined.

Belorusova et al. reported peptides of LXRα comprising the H3 and partial part of H5 correlated to high ABCA1 expression [30]. In addition, the differential relationship between target genes and LXRα isoforms in different patient groups also points to an oncogenic role for LXRα and a tumour suppressor role for LXRβ in TNBC tumours. Indeed, a beneficial role for LXRβ has been reported previously. Both dendrogenin A and RGX-104 are tumour suppressors and selective LXRβ agonists [10, 31, 32].

To the best of our knowledge, a4 (NM_001251934; NP_001238863; B4DXU5), a5 (NM_001363595.2; NP_001350524.1; B5MBY7), and β4 (M0R2F9) have not been reported in the literature before. a4 has an alternative 21 amino acids in the start of the AF1 domain producing a unique peptide mass signature that was detected with MS. These data support its inclusion in UNIPROT with a Q13133 prefix. As this variant has not been investigated before and we perform no functional studies herein, the significance of this partial AF1 substitution remains unknown. α5 was not detected by MS but its presumed protein band (39 kDa) was lost after siLXRα treatment, its transcript was robustly expressed, and protein and transcript expression were strongly correlated. Although a unique peptide for β4 was predicted from *in silico* analyses it was not detected with MS, however, its transcript was robustly expressed, and protein and transcript expression were strongly correlated.

Some of our results are at odds with prior observations. We were unable to detect expression of XP_02430405X (previously termed α4 [16]), in any cell lines or tissues, at RNA or protein level, including in HEPG2 or MCF7 cell lines that were previously shown to express this isoform [16, 33]. Intriguingly, a variant previously reported to be α5 [16, 33] did not correspond to any of the 64 LXRα transcripts from NCBI, nor any LXRα UNIPROT entries **(SF2A).** In this study [16], the measurement of XP_02430405X and previously reported α5 relied on PCR primers that detected a retained intron between exon 6-7 and a retained intron between exon 7-8, respectively **(SF2A).** We went to extensive lengths to ensure primary RNA or gDNA did not contaminate our cDNA libraries, steps that surprisingly have not been reported in LXR splicing studies previously. Cytoplasmic RNA is preferred over total cell isolates when examining splice expression and this has been reported previously to increase sensitivity of splice junction detection [34]. Previous detection of XP_02430405X may therefore be due to amplification of incompletely processed transcripts or contaminating gDNA. During our primer design stage, we found that the PCR primers previously used to measure a1 at the exon 2-3 junction [15, 16, 35] also detect a2 **(SF3).** In addition, primers previously used to measure a2 at the exon 5-7 junction [15, 16, 35] also detect the previously unreported a5 variant (**SF2**). Our correlation analyses in cell lines and primary tumour samples suggest the majority of exon 6 skipping transcripts actually code for a5 not a2. We also found a5 is significantly (20-fold) higher than a2 in MCF7 and equal to a2 in HEPG2, the same cell lines that in prior studies were used to show a2 was strongly expressed. It likely that previous reports have over-estimated the contribution of a2 and under-estimated the contribution of a5 to the LXR pool.

In summary, this study provides critical insight into the pathophysiology of LXR splicing in BCa. Our study clarifies the relative roles of the LXRα and LXRβ isoforms and our data are consistent with the hypothesis that while full length LXRα is oncogenic, LXRβ and truncated LXRα variant (a5) can act as tumour suppressors. We propose the existence of two new LXRα isoforms: a4 containing an alternative AF1 region, and a5 which lacks large sections of the AF1 and LBD, probably rendering it with dominant negative function.

Triple negative breast cancer is defined as a category of exclusion. The data presented here suggest that measuring LXRα1, LXRα5, and LXRβ at RNA or protein level may allow sub-stratification of TNBC patients for increased accuracy of prognosis. Perhaps more importantly, given the array of pharmacological and dietary modulators of the LXR pathway, defining gene networks regulated by the opposing oncogenic and tumour suppressor LXR variants will aid in the development of lifestyle and pharmacological interventions to reduce incidence and improve survival in this challenging cancer of unmet clinical need.

## Acknowledgments

The mass spectrometry was performed and analysis supported by Dr James Ault and Dr Rachel George at the Biomolecular Mass Spectrometry Facility, Faculty of Biological Sciences, University of Leeds.

