## Supplementary Information for "Comprehensive profiling of liver x receptor splicing in triple negative breast cancer reveals existence of novel splice variants that are prognostic for survival"

### Supplementary Materials

Alternative splicing that disrupts ligand binding domains in liver x receptors predicts survival in triple negative breast cancer

Priscilia Lianto[1], J. Bernadette Moore[1], Thomas A. Hughes[2], and James L. Thorne[1].
1School of Food Science and Nutrition, University of Leeds, Leeds, LS2 9JT, UK.
2School of Medicine, University of Leeds, Leeds, LS9 7TF, UK.

#### Supplementary Information Section 1: Isoform naming rationale

Previous publications have used a consistent naming system for the LXRα isoforms, but deviates from nomenclature of NCBI, TCGA Splicing (TSVdb), ENSEMBL, and UNIPROT databases. Due to the complexity of LXR transcript variants [1] the naming system of previous publications was not appropriate [2-4]. For example, NM_001130101 previously referred to as α3 [2-4], is annotated as α2 in the NCBI database. NM_001130102 has previously been referred to as α2 [2, 4], but is annotated as α3 in the NCBI database. Given the large number of transcripts we worked with we adopted the NCBI annotation of the major isoforms. The details of all previous reported LXR splice variants and how they pertain to the major databases is summarised in Table-S1+S2 and transcript and protein structures are summarised in schematic diagrams in **SF1**. During this annotation and documentation process some NCBI predicted LXR isoforms were found to be identical to curated isoforms. For example, the LXRαx5 amino acid sequence is derived from two NCBI predicted isoforms (XP_011518107.1 and XP_005252763.1) and is identical to the LXRα1 amino acid sequence (NP_005684.2). We confirmed this using Clustal Omega Multiple Sequence Alignment software (https://www.ebi.ac.uk/Tools/msa/clustalo/) (EMBL-EBI, Dublin, Ireland). In order to convey the information, we aligned LXRα1 (curated NCBI LXR isoform) to LXRαx5 (predicted NCBI LXR isoform) amino acid sequences as examples of homologous protein isoforms **(SF1C).**

A

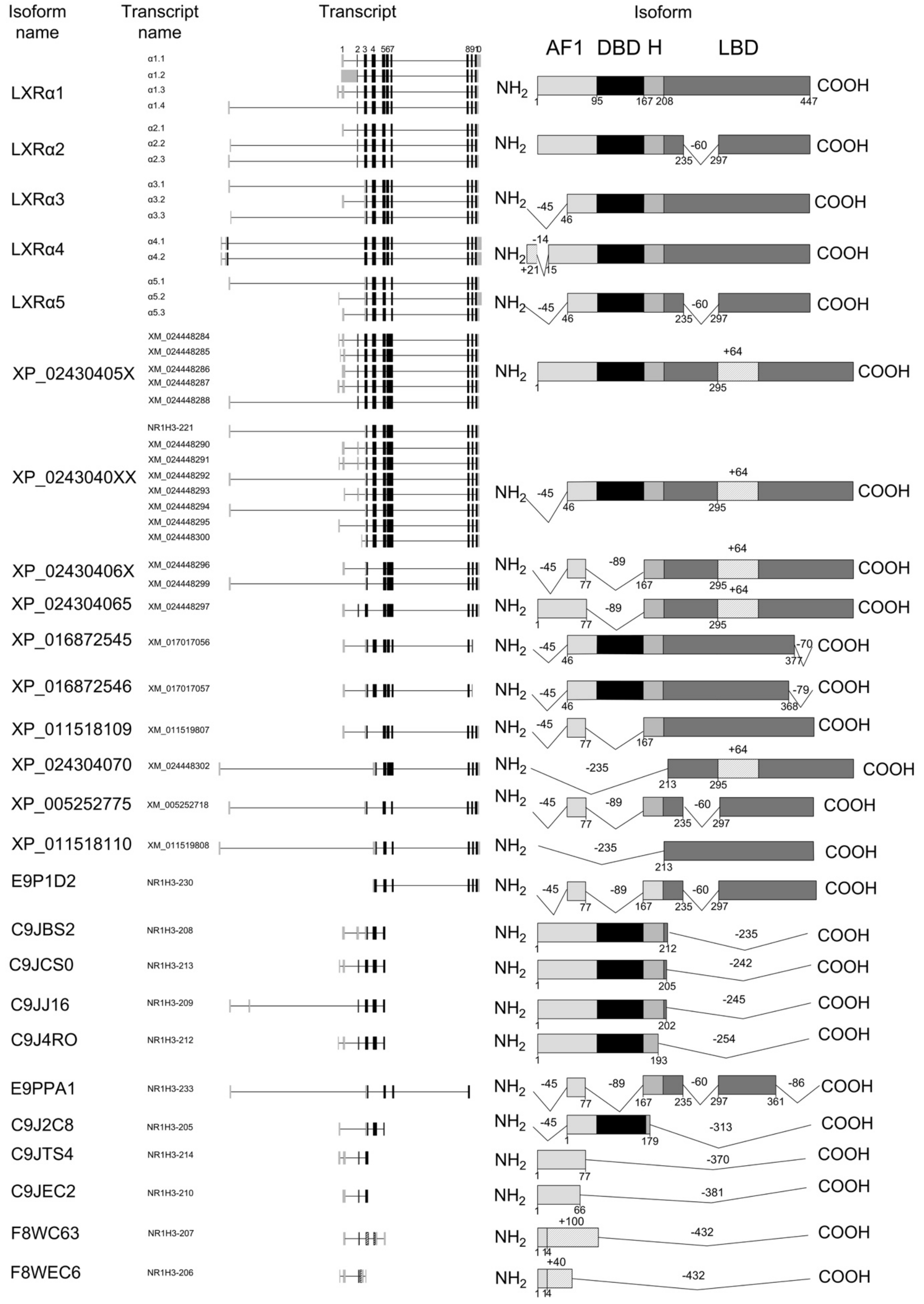

Isoform name
Transcript name
Transcript
Isoform
AF1 DBD H LBD
LXRα1
α1.1
α1.2
α1.3
α1.4
LXRα2
α2.1
α2.2
α2.3
LXRα3
α3.1
α3.2
α3.3
LXRα4
α4.1
α4.2
LXRα5
α5.1
α5.2
α5.3
XP_02430405X
XM_024448284
XM_024448285
XM_024448286
XM_024448287
XM_024448288
XP_0243040XX
NR1H3-221
XM_024448290
XM_024448291
XM_024448292
XM_024448293
XM_024448294
XM_024448295
XM_024448300
XP_02430406X
XM_024448296
XM_024448299
XP_024304065
XM_024448297
XP_016872545
XM_017017056
XP_016872546
XM_017017057
XP_011518109
XM_011519807
XP_024304070
XM_024448302
XP_005252775
XM_005252718
XP_011518110
XM_011519808
E9P1D2
NR1H3-230
C9JBS2
NR1H3-208
C9JCS0
NR1H3-213
C9JJ16
NR1H3-209
C9J4RO
NR1H3-212
E9PPA1
NR1H3-233
C9J2C8
NR1H3-205
C9JTS4
NR1H3-214
C9JEC2
NR1H3-210
F8WC63
NR1H3-207
F8WEC6
NR1H3-206
NH2
COOH

B

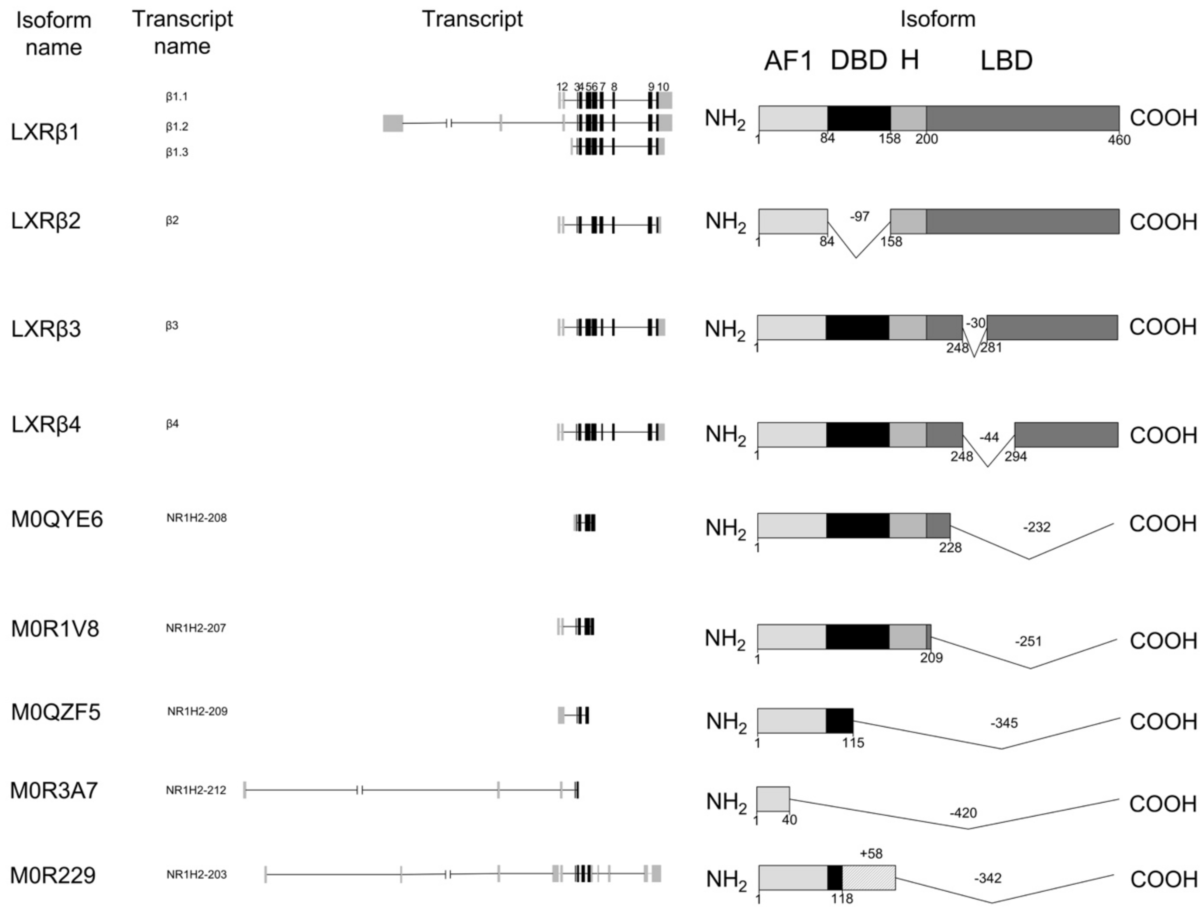

C

```
CLUSTAL O(1.2.4) multiple sequence alignment

α1(NP_005684.2)                        MSLWLGAPVPDIPPDSAVELWKPGAQDASSQAQGGSSCILREEARMPHSAGGTAGVGLEA    60
αx5(XP_011518107.1,XP_005252763.1)     MSLWLGAPVPDIPPDSAVELWKPGAQDASSQAQGGSSCILREEARMPHSAGGTAGVGLEA    60
                                       ************************************************************

α1(NP_005684.2)                        AEPTALLTRAEPPSEPTEIRPQKRKKGPAPKMLGNELCSVCGDKASGFHYNVLSCEGCKG    120
αx5(XP_011518107.1,XP_005252763.1)     AEPTALLTRAEPPSEPTEIRPQKRKKGPAPKMLGNELCSVCGDKASGFHYNVLSCEGCKG    120
                                       ************************************************************

α1(NP_005684.2)                        FFRRSVIKGAHYICHSGGHCPMDTYMRRKCQECRLRKCRQAGMREECVLSEEQIRLKKLK    180
αx5(XP_011518107.1,XP_005252763.1)     FFRRSVIKGAHYICHSGGHCPMDTYMRRKCQECRLRKCRQAGMREECVLSEEQIRLKKLK    180
                                       ************************************************************

α1(NP_005684.2)                        RQEEEQAHATSLPPRASSPPQILPQLSPEQLGMIEKLVAAQQQCNRRSFSDRLRVTPWPM    240
αx5(XP_011518107.1,XP_005252763.1)     RQEEEQAHATSLPPRASSPPQILPQLSPEQLGMIEKLVAAQQQCNRRSFSDRLRVTPWPM    240
                                       ************************************************************

α1(NP_005684.2)                        APDPHSREARQQRFAHFTELAIVSVQEIVDFAKQLPGFLQLSREDQIALLKTSAIEVMLL    300
αx5(XP_011518107.1,XP_005252763.1)     APDPHSREARQQRFAHFTELAIVSVQEIVDFAKQLPGFLQLSREDQIALLKTSAIEVMLL    300
                                       ************************************************************

α1(NP_005684.2)                        ETSRRYNPGSESITFLKDFSYNREDFAKAGLQVEFINPIFEFSRAMNELQLNDAEFALLI    360
αx5(XP_011518107.1,XP_005252763.1)     ETSRRYNPGSESITFLKDFSYNREDFAKAGLQVEFINPIFEFSRAMNELQLNDAEFALLI    360
                                       ************************************************************

α1(NP_005684.2)                        AISIFSADRPNVQDQLQVERLQHTYVEALHAYVSIHHPHDRLMFPRMLMKLVSLRTLSSV    420
αx5(XP_011518107.1,XP_005252763.1)     AISIFSADRPNVQDQLQVERLQHTYVEALHAYVSIHHPHDRLMFPRMLMKLVSLRTLSSV    420
                                       ************************************************************

α1(NP_005684.2)                        HSEQVFALRLQDKKLPPLLSEIWDVHE     447
αx5(XP_011518107.1,XP_005252763.1)     HSEQVFALRLQDKKLPPLLSEIWDVHE     447
                                       ***************************
```

**Supplementary Figure 1. Schematic diagram of LXR splice variants.**

(A) LXRα and (B) LXRβ splice variants reported in NCBI, ENSEMBL, and/or UNIPROT databases. For transcript, black boxes represent translated exons, grey boxes represent untranslated exons, lines joining exons represent intronic regions. The numbers above α1.1 or β1.1 transcripts are referred to the first-discovered exons [5]. For isoform, silver boxes are represented Activation Function 1 (AF1), black boxes are represented DNA Binding Domain (DBD), light grey are represented hinge (H), and dark grey boxes are represented Ligand Binding Domain (LBD). The numbers right below the isoform domain boxes are represented the position of amino acids. The hatched boxes are represented protein domain derived from later-discovered coding exon. The numbers above connecting lines represent missing amino acids. (C) NCBI predicted ($\alpha$1) and curated (ax5) isoforms are fully homologous.

#### Supplementary Information Section 2: Primer design strategy

During primer design, it became clear that due to complexity arising from the large number of LXR coding transcripts, several transcripts were not distinguishable from each other due to the sequence homology and coding redundancy. Thus, some primer pairs detected 'ambiguous amplicons'. This ambiguity was found in several published articles. For example, the PCR primers previously used to measure α1 [2-4] also amplify α2. Primers previously used to measure α2 via the exon 6 skipping (5-7 junction) [2-4] also detect the previously unreported α5 variant. By documenting all possible LXR transcript variants (**SF1, Table S1+S2**), we addressed these issues by a combination of design and experimental approaches and designed unique primer sets against exon-exon junction for most transcript variants.

Primers were designed using NCBI BLAST primer design or, where transcript variants were not available in the NCBI database, primer 3 software was used [6]. According to the gene sequences from NCBI and ENSEMBL, the amplicon size was restricted between 80 to 150 bp. The standard curve from 1-5 ug each concentration cDNA ratio was carried to determine the efficiency of the primers. Primer pairs were chosen based on specificity (single peak in melting temperature analysis) and amplification efficiency. For the purpose of comparison, we summarised primer locations used in previous studies (**SF2A**) and in the current study for LXRα (**SF2B**) and LXRβ (**SF2C**) showing the exon-exon regions in schematic diagrams. All primer sequences used in this study are shown in **Table S3**.

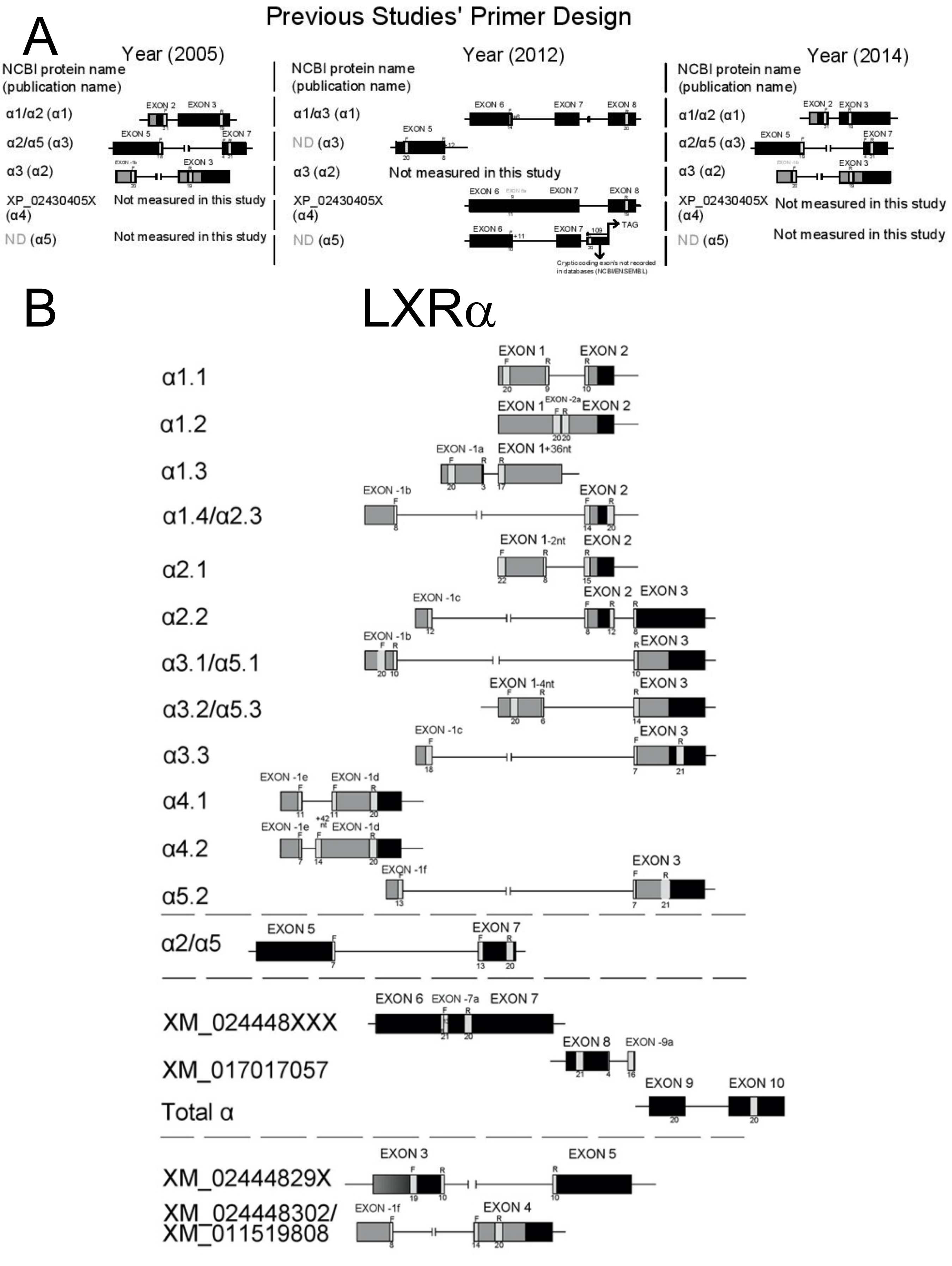
Previous Studies' Primer Design
A
Year (2005)
Year (2012)
Year (2014)
NCBI protein name (publication name)
α1/α2 (α1)
α2/α5 (α3)
α3 (α2)
XP_02430405X (α4)
ND (α5)
α1/α3 (α1)
ND (α3)
Not measured in this study
Cryptic coding exon's not recorded in databases (NCBI/ENSEMBL)
B
LXRα
α1.1
α1.2
α1.3
α1.4/α2.3
α2.1
α2.2
α3.1/α5.1
α3.2/α5.3
α3.3
α4.1
α4.2
α5.2
α2/α5
XM_024448XXX
XM_017017057
Total α
XM_02444829X
XM_024448302/
XM_011519808

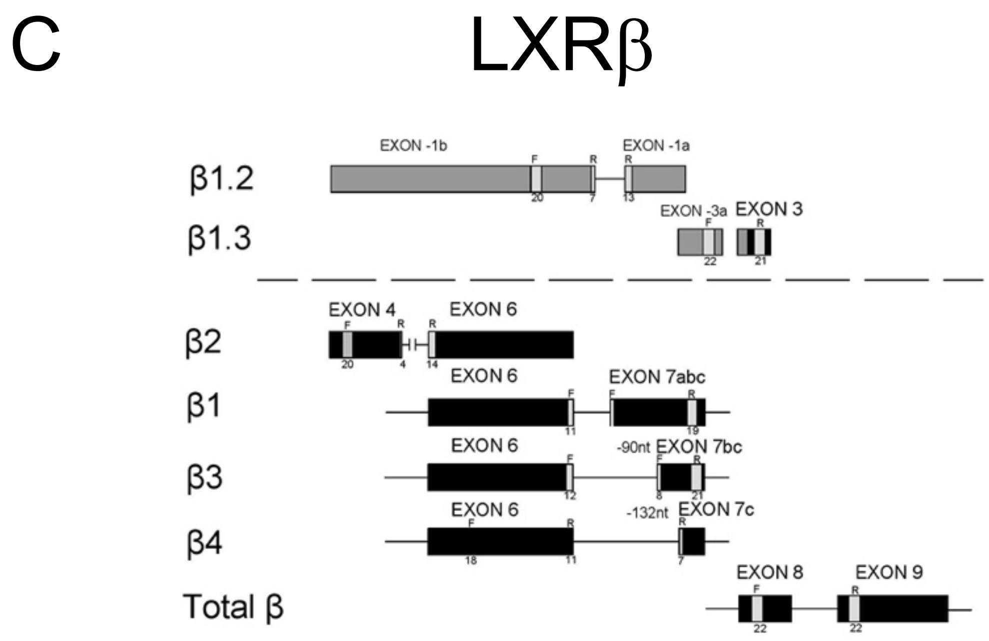

**Supplementary Figure 2: Schematic diagrams illustrating the location of primer pairs for detecting LXR splice variants.**

(A) position of primers and amplicons in previous studies and (B) current study for LXRα and (C) current study for LXRβ. The grey boxes show non-coding exons; the black boxes show coding exons; the lines connecting boxes denote introns; the gray text (e.g., for exon 3 of XM_024448XXX isoforms) indicated that exon 7a can be either coding or non-coding exon. The numeral X in transcript names (e.g. XM_0244829X or XM_024448XXX) denotes multiple variants share these features in NCBI as Ref. Seq. ID. Location of primers is denoted by grey boxes with F (forward) and R (reverse) above and base number below. Primers split between exons have number of bases to which the primers bind on each side of the boundary. ND = transcript not documented in any database.

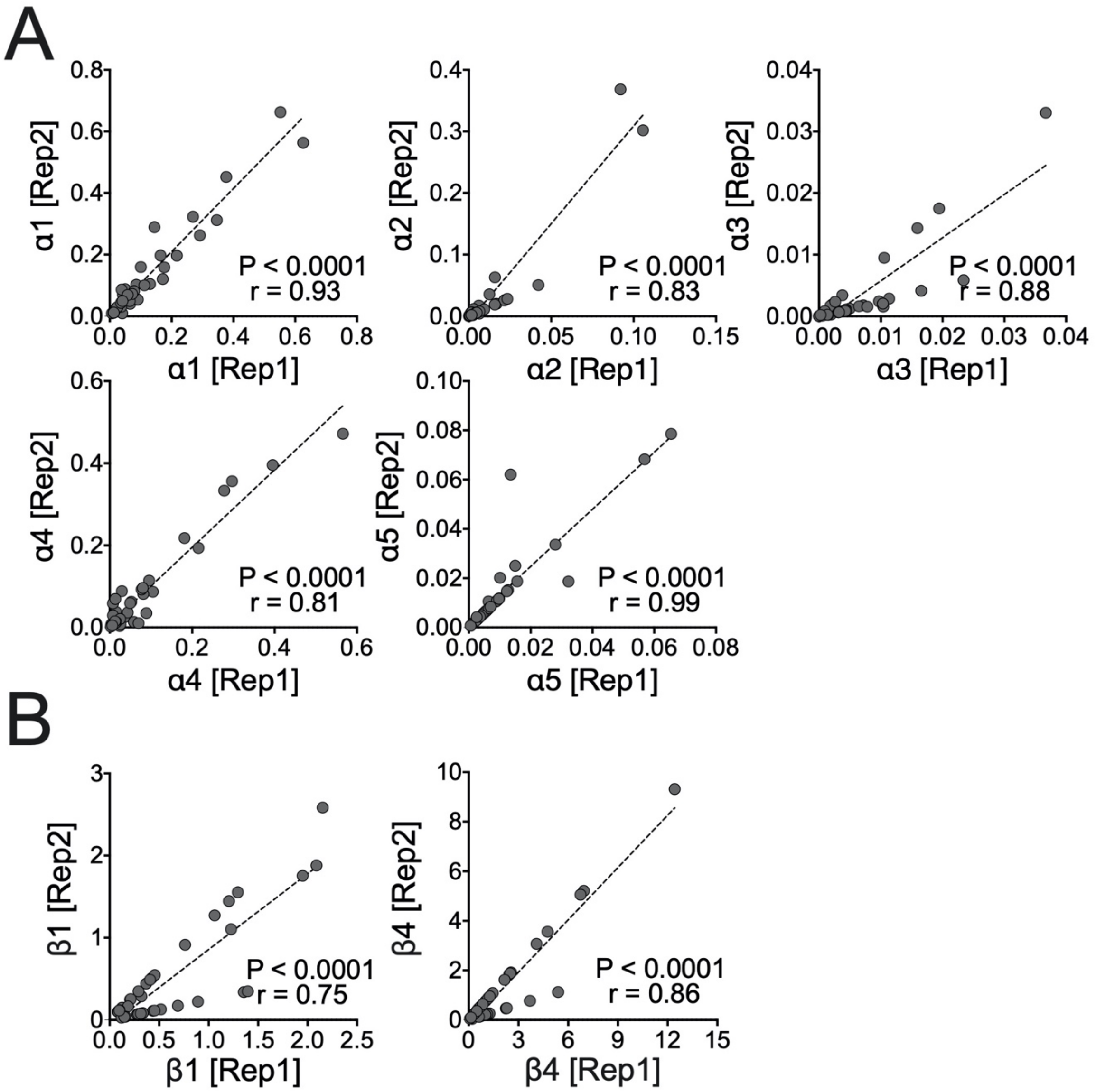

**Supplementary Figure 3. Correlation between two slices of each tumour sample when measuring LXR protein variant expression.**

LXRα (A) or LXRβ (B) protein variant expression of tumour replicate 1 (x axis) is plotted against replicate 2 (y axis). Replicate 1 and 2 were assigned randomly. Samples were derived from the same TNBC tumour sample, normalised by HPRT expression level, and expression correlated using Interclass correlation coefficient through Spearman's rank [7]. Circles represented individual TNBC samples patients. Significance was considered when p≤0.05.

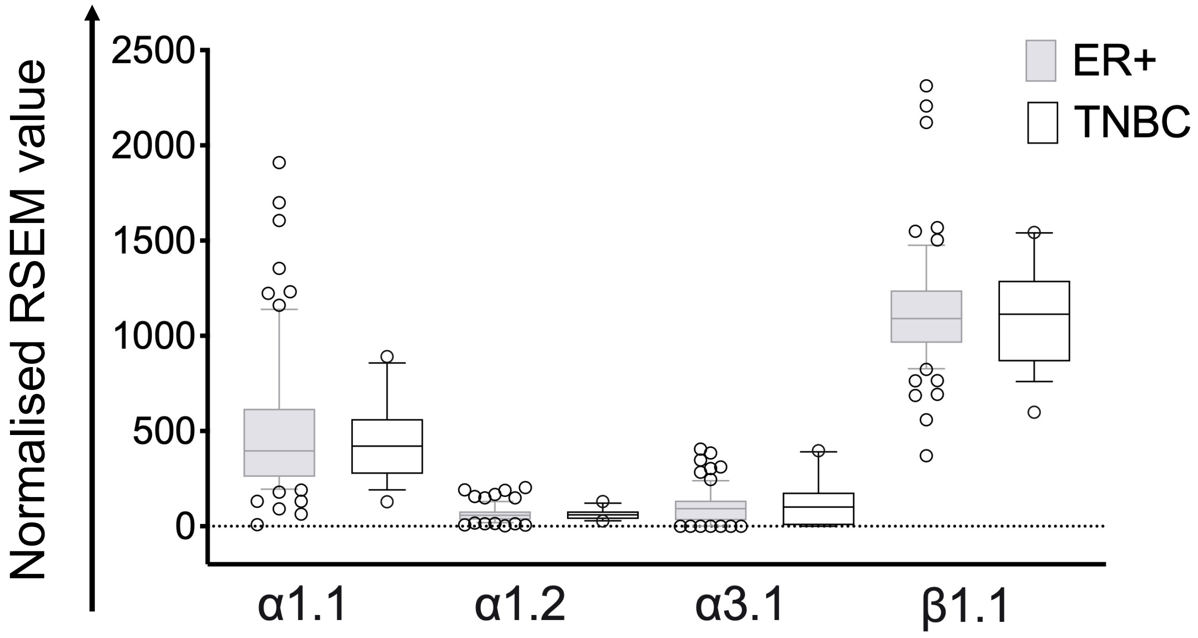

**Supplementary Figure 4. LXR transcript variant expression in ER+ and TNBC tumour samples from the TSVdb TCGA tumour cohort.**

Differential expression of LXR transcript variants in ER+ and TNBC tumour samples were plotted in box and whisker charts. The box extends from the 10th to 90th percentile. The line in the middle of the box shows median. The whiskers show minimum and maximum values. Significance was considered when $p \leq 0.05$.

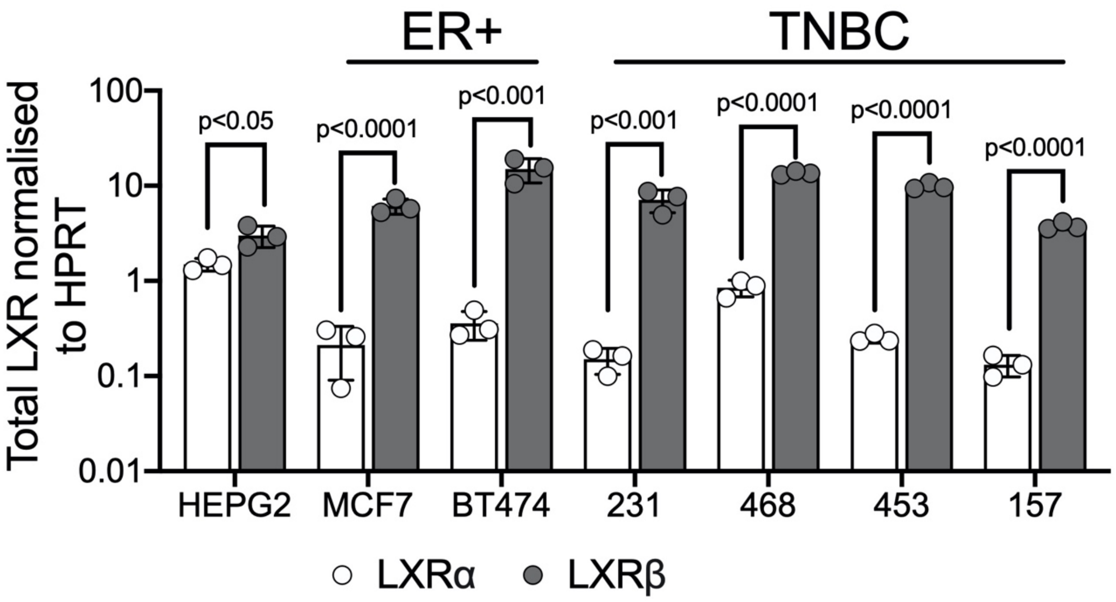

**Supplementary Figure 5. LXRβ is expressed at higher levels than LXRα in all breast cancer cell lines.**

Data showed mean of three independent replicates with SEM. p-values calculated using multiple two-way t-tests. Significance was considered when $p \leq 0.05$.

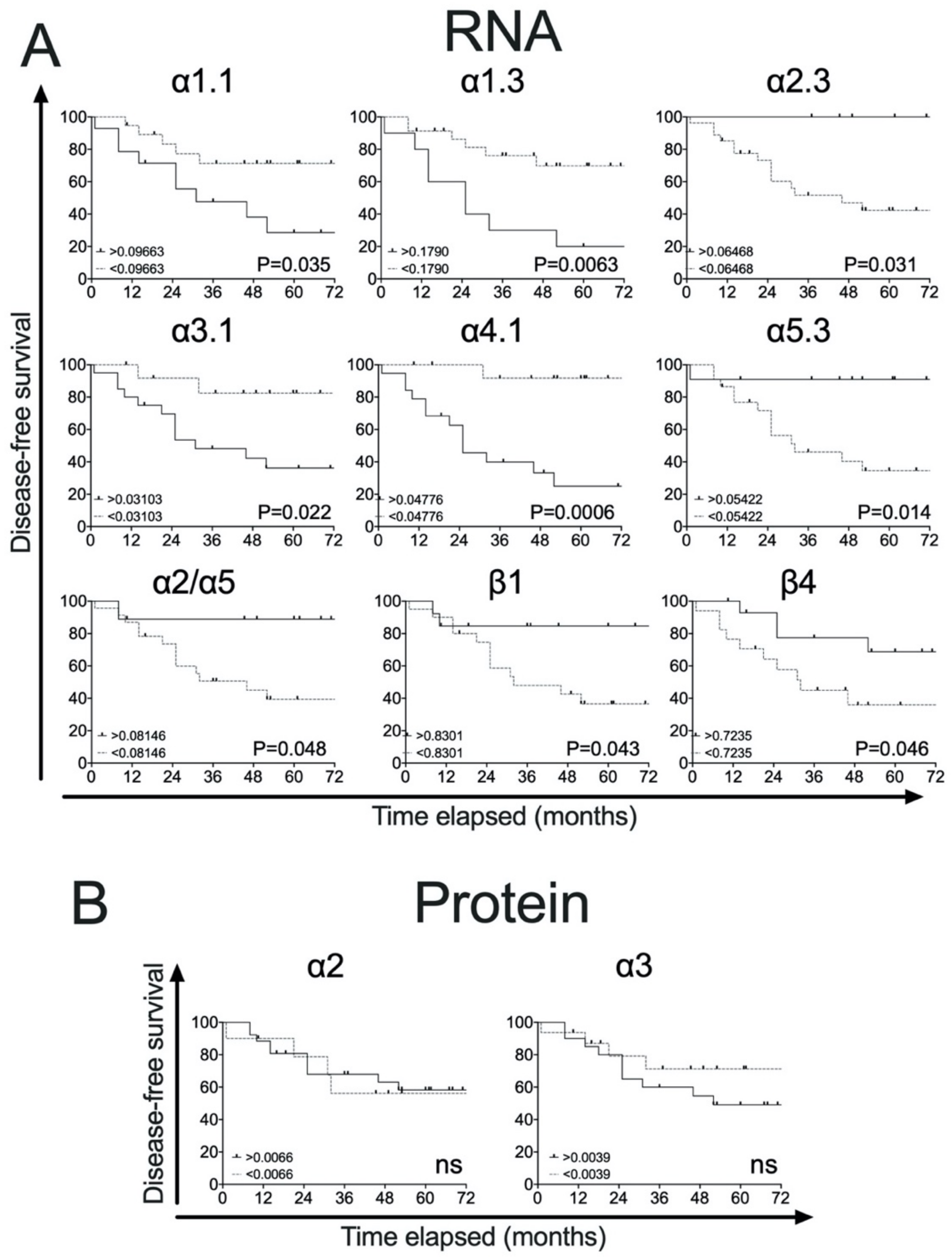

**Supplementary Figure 6. Splice variants and TNBC patient survival.**

(A) All transcript variants are prognostic. (B) LXRα2 and α3 protein variants are not prognostic. TNBC patients were divided into two groups (n=38), no event (n=23) and event (n=15), based on their disease-free survival status. Kaplan-Meier survival curves plotting disease free survival of TNBC patients with high or low variant expression relative to HPRT. Data derived from the mean of two different slices of tumours. Significance determined by the Log-rank (Mantel-cox) test, $p \leq 0.05$ was considered significant.

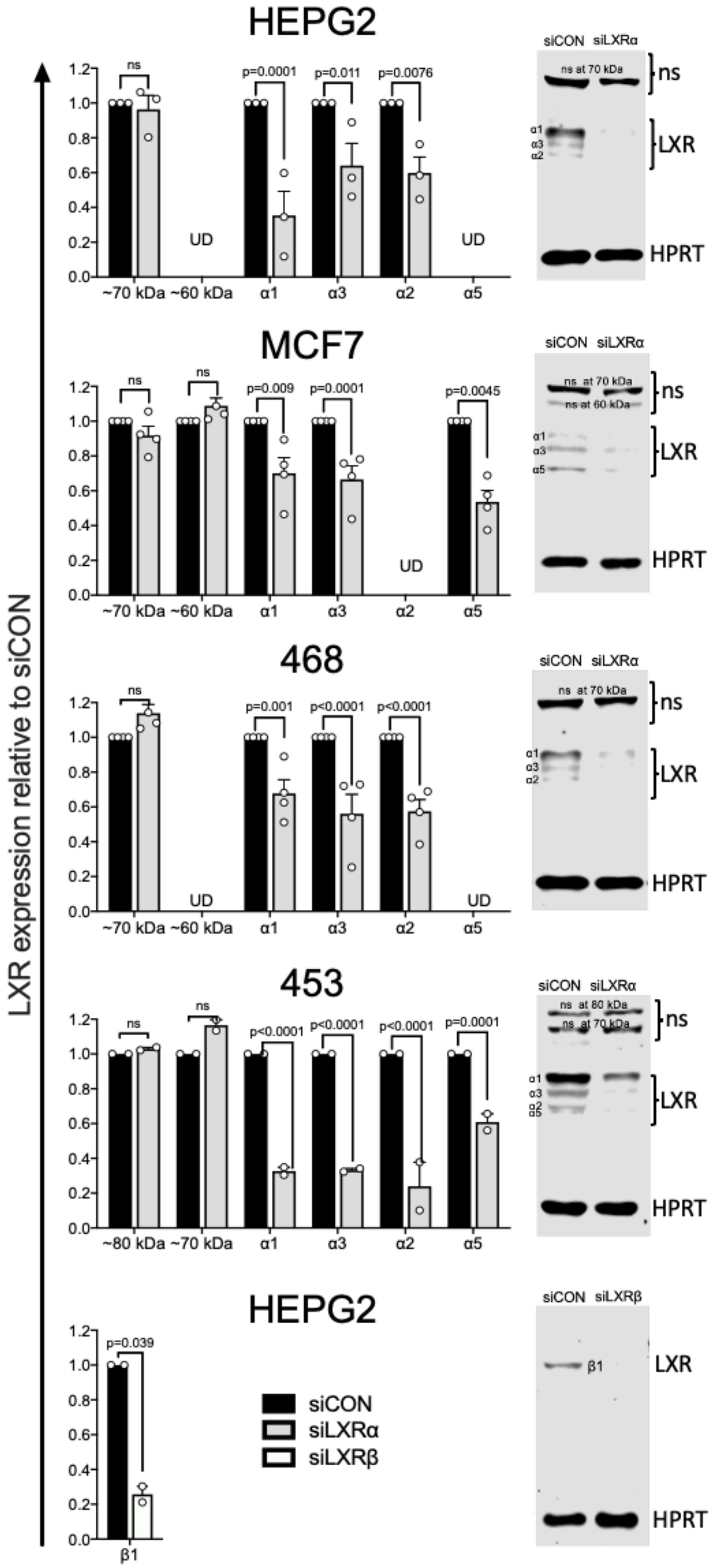
HEPG2
MCF7
468
453
HEPG2
LXR expression relative to siCON
~70 kDa ~60 kDa α1 α3 α2 α5
~80 kDa ~70 kDa α1 α3 α2 α5
β1
ns
p=0.0001
p=0.011
p=0.0076
p=0.009
p=0.0045
p=0.001
p<0.0001
p=0.039
UD
siCON
siLXRα
siLXRβ
ns at 70 kDa
ns at 60 kDa
ns at 80 kDa
LXR
HPRT

**Supplementary Figure 7. Relative expression LXR protein variants to control siRNA after siLXRα or siLXRβ treatment.**

Protein expression determined in ImageJ. Data represent mean of 2-4 independent replicates with SEM. Additional bands at 60kDa and 70kDa were assumed to be non-specific (ns) as their sizes did not correspond to any of the 48 LXRα transcripts. As expected, non-specific bands were unaffected by siRNA treatment. Statistical significance was measured by two-tailed one-way ANOVA, p-values are given above bars showing significant difference compared to siCON. siCON = siControl.

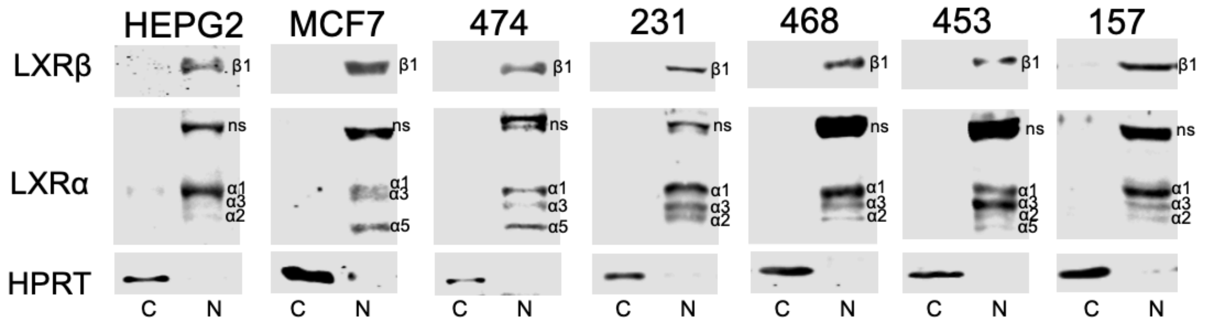

**Supplementary Figure 8. All LXR splice variants are localized in nucleus.**

HPRT is used as cytoplasmic control. ns = non-specific binding; C=cytoplasmic; N=nuclear.

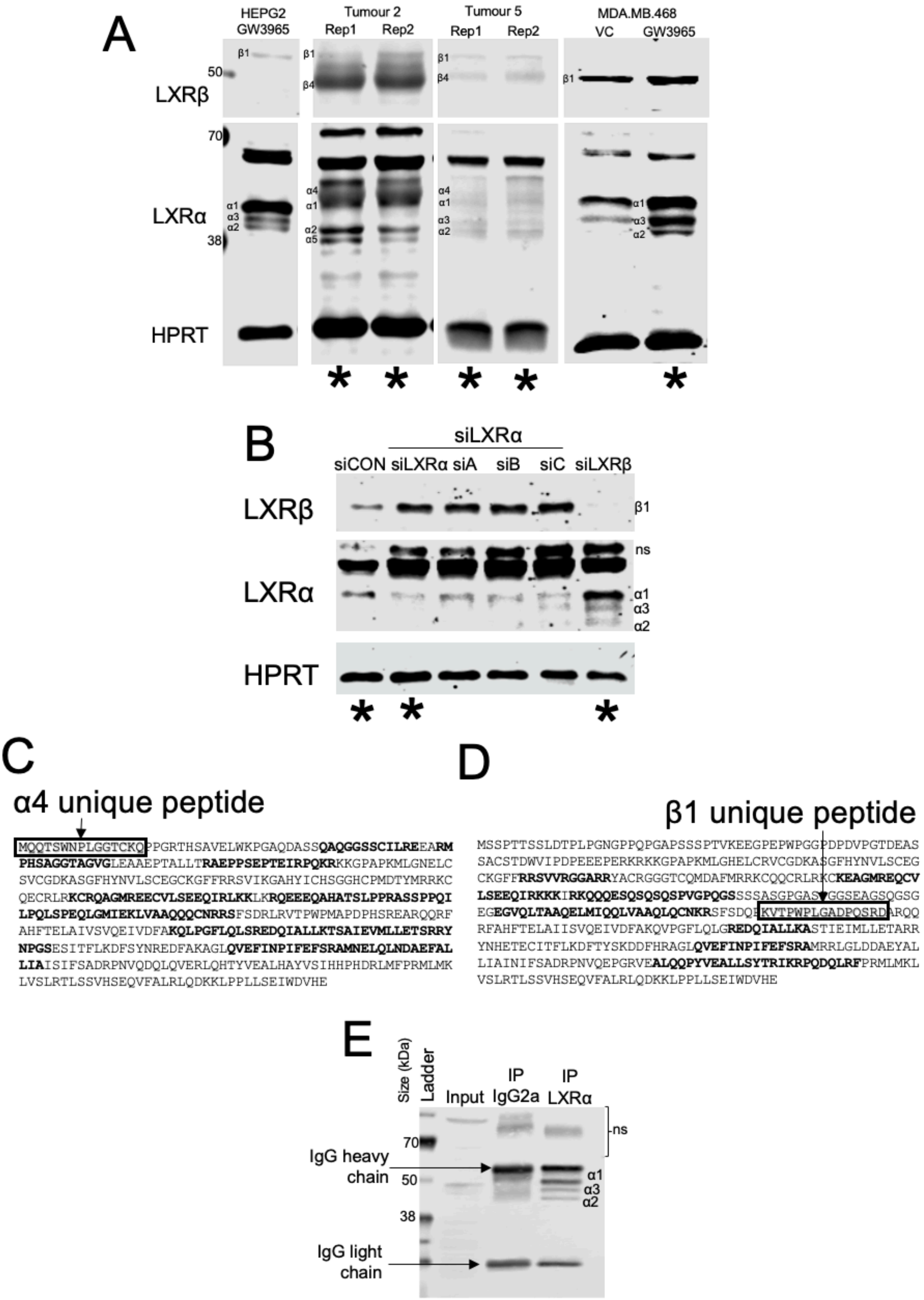
A
HEPG2
GW3965
Tumour 2
Rep1
Rep2
Tumour 5
Rep1
Rep2
MDA.MB.468
VC
GW3965
LXRβ
LXRα
HPRT
50
70
38
β1
β4
α1
α2
α3
α4
α5
*
B
siLXRα
siCON
siLXRα
siA
siB
siC
siLXRβ
LXRβ
LXRα
HPRT
β1
ns
α1
α3
α2
*
C
α4 unique peptide
MQQTSWNPLGGTCKQPPGRTHSAVELWKPGAQDASSQAQGGSSCILREEARM
PHSAGGTAGVGLEAAEPTALLTRAEPPSEPTEIRPQKRKKGPAPKMLGNELC
SVCGDKASGFHYNVLSCEGCKGFFRRSVIKGAHYICHSGGHCPMDTYMRRKC
QECRLRKCRQAGMREECVLSEEQIRLKKLKRQEEEQAHATSLPPRASSPPQI
LPQLSPEQLGMIEKLVAAQQQCNRRSFSDRLRVTPWPMAPDPHSREARQQRF
AHFTELAIVSVQEIVDFAKQLPGFLQLSREDQIALLKTSAIEVMLLETSRRY
NPGSESITFLKDFSYNREDFAKAGLQVEFINPIFEFSRAMNELQLNDAEFAL
LIAISIFSADRPNVQDQLQVERLQHTYVEALHAYVSIHHPHDRLMFPRMLMK
LVSLRTLSSVHSEQVFALRLQDKKLPPLLSEIWDVHE
D
β1 unique peptide
MSSPTTSSLDTPLPGNGPPQPGAPSSSPTVKEEGPEPWPGGPDPDVPGTDEAS
SACSTDWVIPDPEEEPERKRKKGPAPKMLGHELCRVCGDKASGFHYNVLSCEG
CKGFFRRSVVRGGARRYACRGGGTCQMDAFMRRKCQQCRLRKCKEAGMREQCV
LSEEQIRKKKIRKQQQESQSQSQSPVGPQGSSSSASGPGASPGGSEAGSQGSG
EGEGVQLTAAQELMIQQLVAAQLQCNKRSFSDQPKVTPWPLGADPQSRDARQQ
RFAHFTELAIISVQEIVDFAKQVPGFLQLGREDQIALLKASTIEIMLLETARR
YNHETECITFLKDFTYSKDDFHRAGLQVEFINPIFEFSRAMRRLGLDDAEYAL
LIAINIFSADRPNVQEPGRVEALQQPYVEALLSYTRIKRPQDQLRFPRMLMKL
VSLRTLSSVHSEQVFALRLQDKKLPPLLSEIWDVHE
E
Size (kDa)
Ladder
Input
IP
IgG2a
IP
LXRα
70
50
38
ns
IgG heavy
chain
IgG light
chain
α1
α3
α2

**Supplementary figure 9. Representative blots showing the LXR splicing protein expression in samples subjected to Mass-Spectrometry (MS).**

(A) TNBC tumours and in MDA.MB.468 cell line samples treated with GW3965, siLXRα or siLXRβ treated MDA.MB.468 cells. Samples marked with asterix were subjected to mass spectrometry (MS) analysis. Sequence coverage of unique (boxes) or homologous (bold) peptides LXRα (C) and LXRβ (D) peptides identified by proteomic analysis. Homologous peptides are generated by multiple LXR splice variants and can’t be used to confirm the presence or individual variants. (E) MDA.MB.468 subjected to immunoprecipitation. VC = vehicle control; T = tumour; siA = siExon4 of LXRα, siB = siExon9-10 (coding exon) of LXRα, siC = siExon 10 (non-coding 3’UTR) of LXRα, siLXRα = siA+siB+siC. ns = non-specific binding.

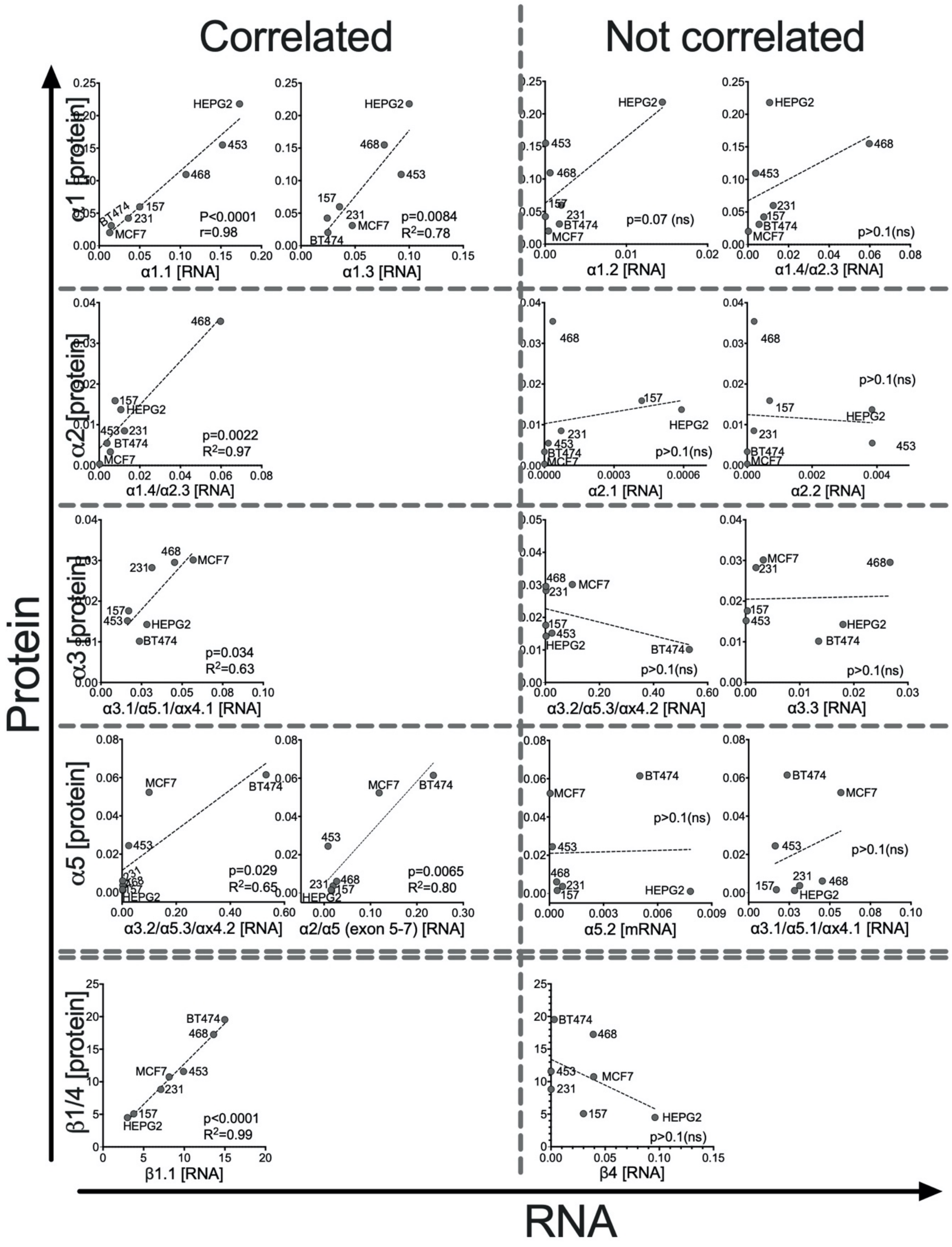

**Supplementary figure 10. Correlation of LXR transcripts with LXR protein variants in cell lines.**

RNA and protein normalised to HPRT levels, and significance tested for using linear regression. Circles represent mean of 3-5 separate passage of each cell line. P-values and R-squared ($R^2$) are shown in each graph. NS = not significant.

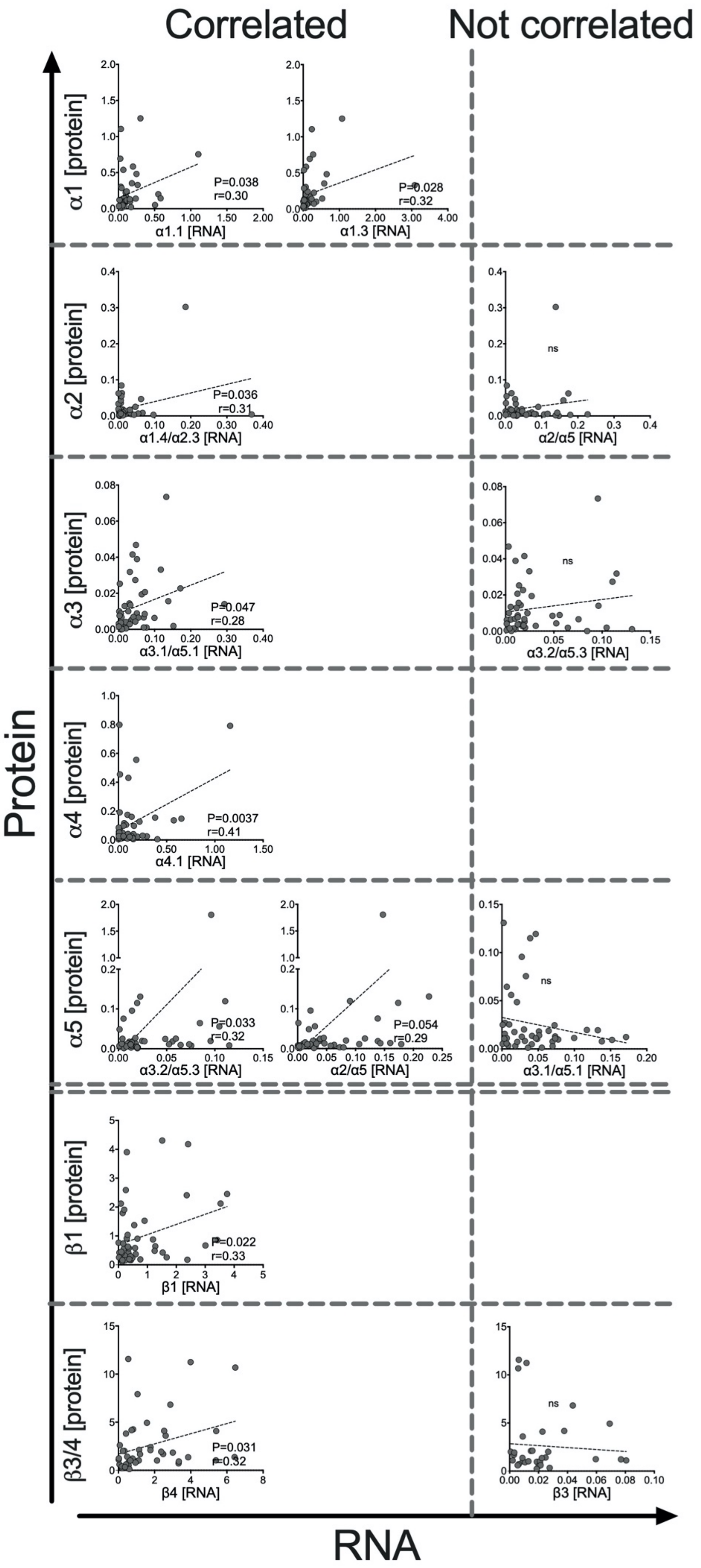

Correlated
Not correlated
Protein
RNA
α1 [protein]
α1.1 [RNA]
P=0.038
r=0.30
α1.3 [RNA]
P=0.028
r=0.32
α2 [protein]
α1.4/α2.3 [RNA]
P=0.036
r=0.31
α2/α5 [RNA]
ns
α3 [protein]
α3.1/α5.1 [RNA]
P=0.047
r=0.28
α3.2/α5.3 [RNA]
ns
α4 [protein]
α4.1 [RNA]
P=0.0037
r=0.41
α5 [protein]
α3.2/α5.3 [RNA]
P=0.033
r=0.32
α2/α5 [RNA]
P=0.054
r=0.29
α3.1/α5.1 [RNA]
ns
β1 [protein]
β1 [RNA]
P=0.022
r=0.33
β3/4 [protein]
β4 [RNA]
P=0.031
r=0.32
β3 [RNA]
ns

**Supplementary figure 11. Correlation of LXR transcripts with LXR protein variants in TNBC tumours.**

RNA and protein normalised to HPRT levels, and significance tested for using linear regression. Circles represent mean of two different tumour slices from one TNBC tumour samples. P-values and R-squared ($R^2$) are shown in each graph. NS = not significant.

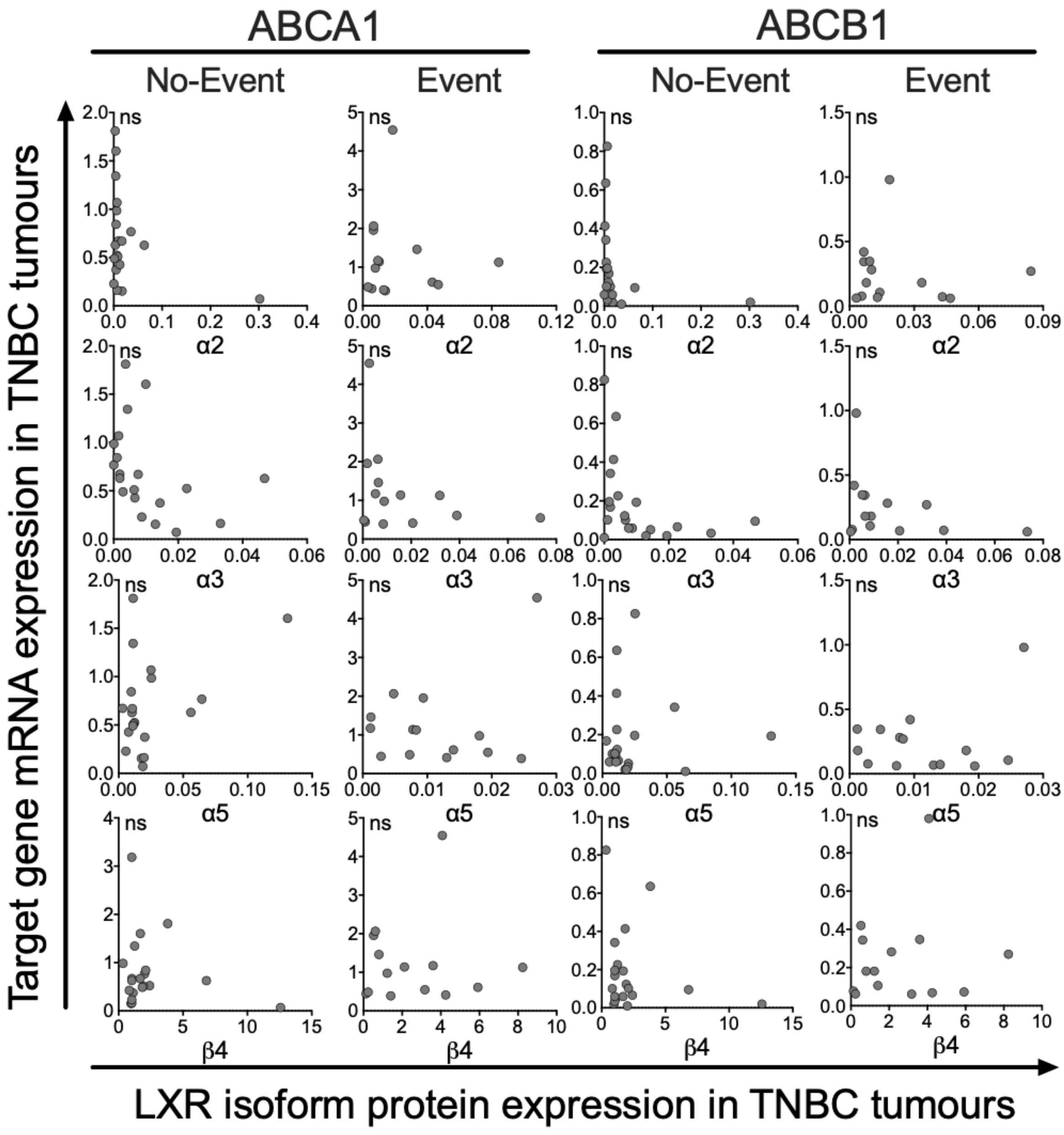

**Supplementary Figure 12. LXRα2, α3, α5, and β4 protein were not correlated to target genes.**

TNBC patients (n=38) were divided into two groups, no event (n=23) and event (n=15), based on their disease-free survival status. LXRα or LXRβ protein variants at x axis versus (A) ABCA1 or (B) ABCB1 gene expression levels at y axis in TNBC tumours were normalised for HPRT levels and correlated using linear regression. Circles represent individual TNBC samples from the mean of two different tumour slices. Significance levels were set at p-values ≤ 0.05.

#### Table S1. The summary information of the different name used for LXRα spice variants

| This study: protein designation | NCBI protein designation | This study: transcript designation | NCBI nucleotide Ref. Seq. Hyperlinked | NCBI protein Ref. Seq. Hyperlinked | ENSEMBL ID | TSVdb ID | UNIPROT ID | Year of publication (doi): hyperlinked | Previous designation | Total amino acids | Size (kDa) | Notes |
|---|---|---|---|---|---|---|---|---|---|---|---|---|
| LXRα1 | LXRα isoform 1 | α1.1 | NM_005693.4 | NP_005684.2 | NR1H3-211 | uc009ylm | Q13133-1 | 1995 (10.1101/gad.9.9.1033);<br>2005 (10.1194/jlr.M500157-JLR200);<br>2012 (10.1124/mol.111.077206); 2014 (10.1016/j.fertnstert.2014.04.033) | LXRα1 | 447 | 50.41 | Detected by PCR and immunoblotting<br>X5 shares 100% amino acid homology |
| | | α1.2 | | | NR1H3-217 | uc001nem | | | | | | |
| | LXRα isoform X5* | α1.3 | XM_005252706.1 | XP_005252763.1 | | | | | | | | |
| | LXRα isoform X5* | α1.4 | XM_011519805.2 | XP_011518107.1 | | | | | | | | |
| LXRα2 | LXRα isoform 2 | α2.1 | NM_001130101.3 | NP_001123573.1 | NR1H3-203 | uc001nen | Q13133-2 | 2005 (10.1194/jlr.M500157-JLR200);<br>2012 (10.1124/mol.111.077206); 2014 (10.1016/j.fertnstert.2014.04.033) | LXRα3 | 387 | 43.56 | Detected by PCR and immunoblotting<br>X11 shares 100% amino acid homology |
| | | α2.2 | | | NR1H3-204 | | | | | | | |
| | LXRα isoform X11* | α2.3 | XM_005252713.3 | XP_005252770.1 | | | | | | | | |
| LXRα3 | LXRα isoform 3 | α3.1 | NM_001130102.3 | NP_001123574.1 | NR1H3-201 | uc001nek | Q13133-3 | 2005 (10.1194/jlr.M500157-JLR200);<br>2012 (10.1124/mol.111.077206); 2014 (10.1016/j.fertnstert.2014.04.033) | LXRα2 | 402 | 45.69 | Detected by PCR and immunoblotting<br>X2 shares 100% amino acid homology |
| | LXRα isoform X2* | α3.2 | XM_024448289.1 | XP_024304057.1 | | | | | | | | |
| | LXRα isoform X2* | α3.3 | XM_024448298.1 | XP_024304066.1 | | | | | | | | |
| LXRα4 | LXRα isoform 4 | α4.1 | NM_001251934.1 | NP_001238863.1 | NR1H3-235 | uc010rhk | B4DXU5 | | | 435 | 51.11 | Detected by PCR, immunoblotting and MS by unique peptide |
| | LXRα isoform 4 | α4.2 | NM_001251935.1 | NP_001238864.1 | | uc009yll | | | | | | |
| LXRα5 | LXRα isoform 5 | α5.1<br>α5.2 | NM_001363595.2 | NP_001350524.1 | NR1H3-202 | | B5MBY7 | | | 342 | 38.85 | Detected by PCR and immunoblotting<br>X7 shares 100% amino acid homology |
| | LXRα isoform X7* | α5.3 | XM_011519806.1 | XP_011518108.1 | | | | | | | | |
| XP_02430405X | LXRα isoform X1 | XM_02444828X | XM_024448284.1<br>XM_024448285.1<br>XM_024448286.1<br>XM_024448287.1<br>XM_024448288.1 | XP_024304052.1<br>XP_024304053.1<br>XP_024304054.1<br>XP_024304055.1<br>XP_024304056.1 | | | | 2012 (10.1124/mol.111.077206) | LXRα4 | 511 | 57.53 | Not detected |
| XP_0243040XX | LXRα isoform X3 | XM_024448XXX | XM_024448290.1<br>XM_024448291.1<br>XM_024448292.1<br>XM_024448293.1<br>XM_024448294.1<br>XM_024448295.1<br>XM_024448300.1 | XP_024304058.1<br>XP_024304059.1<br>XP_024304060.1<br>XP_024304061.1<br>XP_024304062.1<br>XP_024304063.1<br>XP_024304068.1 | NR1H3-221 | | E9PLL4 | | | 466 | 52.81 | Not detected |
| XP_02430406X | LXRα isoform X4 | XM_02444829X | XM_024448296.1<br>XM_024448299.1 | XP_024304064.1<br>XP_024304067.1 | | | | | | 377 | 42.64 | Not detected |
| XP_024304065.1 | LXRα isoform X6 | XM_024448297.1 | XM_024448297.1 | XP_024304065.1 | | | | | | 422 | 47.34 | Not detected |
| XP_016872545.1 | LXRα isoform X8 | XM_017017056.1 | XM_017017056.1 | XP_016872545.1 | | | | | | 332 | 37.25 | Not measured (no unique exon-exon boundaries) |
| XP_016872546.1 | LXRα isoform X9 | XM_017017057.1 | XM_017017057.1 | XP_016872546.1 | | | | | | 323 | 36.26 | Detected by PCR |
| XP_011518109.1 | LXRα isoform X10 | XM_011519807.1 | XM_011519807.1 | XP_011518109.1 | | | | | | 313 | 35.52 | Not measured (no unique exon-exon boundaries) |
| XP_024304070.1 | LXRα isoform X12 | XM_024448302.1 | XM_024448302.1 | XP_024304070.1 | | | | | | 299 | 34.44 | Detected by PCR |
| XP_005252775.1 | LXRα isoform X13 | XM_005252718.3 | XM_005252718.3 | XP_005252775.1 | | | | | | 253 | 28.68 | Not measured (no unique exon-exon boundaries) |
| XP_011518110.1 | LXRα isoform X14 | XM_011519808.2 | XM_011519808.2 | XP_011518110.1 | | | | | | 235 | 27.32 | Detected by PCR |

| | | | | | | |
|---|---|---|---|---|---|---|
| E9P1D2 | NR1H3-230 | NR1H3-230 | E9P1D2 | 296 | 34.14 | Not measured (no unique exon-exon boundaries) |
| C9JBS2 | NR1H3-208 | NR1H3-208 | C9JBS2 | 212 | 23.10 | Not measured (no unique exon-exon boundaries) |
| C9JCS0 | NR1H3-213 | NR1H3-213 | C9JCS0 | 205 | 22.38 | Not measured (no unique exon-exon boundaries) |
| C9JJ16 | NR1H3-209 | NR1H3-209 | C9JJ16 | 202 | 22.04 | Not measured (no unique exon-exon boundaries) |
| C9J4RO | NR1H3-212 | NR1H3-212 | C9J4RO | 193 | 21.10 | Not measured (no unique exon-exon boundaries) |
| E9PPA1 | NR1H3-233 | NR1H3-233 | E9PPA1 | 167 | 18.55 | Not measured (no unique exon-exon boundaries) |
| C9J2C8 | NR1H3-205 | NR1H3-205 | C9J2C8 | 134 | 14.79 | Not measured (no unique exon-exon boundaries) |
| C9JTS4 | NR1H3-214 | NR1H3-214 | C9JTS4 | 77 | 7.82 | Not measured |
| C9JEC2 | NR1H3-210 | NR1H3-210 | C9JEC2 | 68 | 6.86 | Not measured |
| F8WC63 | NR1H3-207 | NR1H3-207 | F8WC63 | 114 | 12.29 | Not measured (non-sense mediated decay) |
| F8WEC6 | NR1H3-206 | NR1H3-206 | F8WEC6 | 54 | 5.71 | Not measured (non-sense mediated decay) |
| | | NR1H3-215<br>NR1H3-218<br>NR1H3-219<br>NR1H3-226<br>NR1H3-228<br>NR1H3-229<br>NR1H3-231<br>NR1H3-232<br>NR1H3-234<br>NR1H3-216<br>NR1H3-220<br>NR1H3-222<br>NR1H3-223<br>NR1H3-224<br>NR1H3-225<br>NR1H3-227 | Processed transcripts [no protein produced] | | | |

### Table S2. The summary information of the different name used for LXRβ spice variants

| **This study: protein designation** | NCBI protein designation | This study: transcript designation | NCBI nucleotide Ref. Seq. Hyperlinked | NCBI protein Ref. Seq. Hyperlinked | ENSEMBL ID | TSVdb ID | UNIPROT ID | Year of publication (doi): hyperlinked | Previously published designation | Total amino acids | Size (kDa) | Notes |
|---|---|---|---|---|---|---|---|---|---|---|---|---|
| **LXRβ1** | LXRβ isoform 1 | **β1.1** | NM_007121.7 | NP_009052.4 | NR1H2-201 | uc010enw | P55055-1 | 1995(10.1101/gad.9.9.1033); 2014(10.1016/j.fertnstert.2014.04.033) | LXRβ | 460 | 50.97 | Detected by PCR and immunoblotting |
| | | **β1.2** | | | NR1H2-214 | | | | | | | Not detected |
| | | **β1.3** | XM_005252706.1 | XP_005252763.1 | NR1H2-204 | | | | | | | |
| **LXRβ2** | LXRβ isoform 2 | **β2** | NM_001256647.3 | NP_001243576.2 | NR1H2-202 | uc002psa | P55055-2 | | | 363 | 39.92 | Not detected |
| **LXRβ3** | | **β3** | | | NR1H2-210 | | M0R0K3 | | | 430 | 47.56 | Detected by PCR |
| **LXRβ4** | | **β4** | | | NR1H2-211 | | M0R2F9 | | | 416 | 45.98 | Detected by PCR and immunoblotting |
| **M0QYE6** | | **NR1H2-208** | | | NR1H2-208 | | M0QYE6 | | | 228 | 24.13 | Not measured (no unique exon-exon boundaries) |
| **M0R1V8** | | **NR1H2-207** | | | NR1H2-207 | | M0R1V8 | | | 209 | 22.24 | Not measured (no unique exon-exon boundaries) |
| **M0QZF5** | | **NR1H2-209** | | | NR1H2-209 | | M0QZF5 | | | 115 | 12.13 | Not measured (no unique exon-exon boundaries) |
| **M0R3A7** | | **NR1H2-212** | | | NR1H2-212 | | M0R3A7 | | | 40 | 3.97 | Not measured (no unique exon-exon boundaries) |
| **M0R229 (non-sense mediated decay)** | | **NR1H2-203** | | | NR1H2-203 | | M0R229 (non-sense mediated decay) | | | 176 | 18.73 | Not measured (non-sense mediated decay) |
| | | | | | NR1H2-213<br>NR1H2-206<br>NR1H2-205 | | Processed transcripts [no protein produced] | | | | | |

**Table S3. TNBC patient tumour characteristics**

| Characteristic | Category | Leeds Breast Research Tissue Bank No. of patients = 38 (%) |
|---|---|---|
| Invasive Tumour grade | 1 | 0 (0) |
| | 2 | 3 (8) |
| | 3 | 35 (92) |
| Tumour size | ≤35 mm | 29 (76) |
| | >35 mm | 9 (24) |
| Survival status | Alive | 28 (74) |
| | Deceased | 10 (26) |
| Recurrence/metastasis | None | 23 (60) |
| | Local and/or distal | 15 (40) |

**Table S4. Primer Sequences of LXR transcripts for qPCR**

| LXR transcript | Forward | Reverse |
|---|---|---|
| α1.1 | TGCTCAGCTCCAGCTCACTG | AGGCACTGTCCAAATCCCCA |
| α1.2 | TCTGGGGAGAAGTGAGGGGT | CACTTTCCAGGGTCCCAGCA |
| a1.3 | CCTATGGAGGGGAGGGAACA | TGAGCACAAGCAGGACCCAG |
| α1.4/α2.3 | AGGAGCATAAGAAGGACAGTGC | GAGGAATGTCAGGCACAGGG |
| α2.1 | CTCAGCCTTTCCCCAAATTGCT | TACCAAGGCACTGTCCAAATCC |
| α2.2 | GAGCATAAGAAG-GACAGTGC | CGCAGAGTCAGGAGGAATGT |
| α3.1/α5.1* | GAAAAGGCGCAGTCTCGGTG | ACCGCAGAGTCTTCTTATGCT |
| α3.2/α5.3* | CACCGAGACTTCTGGACAGG | CCACCGCAGAGTCAAATCCC |
| α3.3 | GGAGAAATCCCTTACCAAGACTCTG | GCATCCTGGCTTCCTCTCTGA |
| α4.1 | GCCAAGGTACAGGTAACGAAGC | TCCTTCTCGGCGTGAACCTG |
| α4.2 | AGGTACAGCTTCAGGGAAGTC | CTCCTTCTCGGCGTGAACCT |
| α5.2 | GCTAAGAGCGCTGGACTCTG | GCATCCTGGCTTCCTCTCTGA |
| α2/α5* | GAGTCACGGTGATGCTTCTG | TGGCAAAGTCTTCCCGGTTA |
| XM_024448XXX | GATCGAGGTGGCTGGAGAAG | AACCTCAAACGGGGACTAGG |
| XM_02444829X | CACTCTGCTGGGGGTACTG | GACAGGACACCTGTGGGTTC |
| XM_017017057 | CATCTTCGAGTTCTCCAGGGC | GCCAAGCTCTCTCATCCTGC |
| XM_024448302/ XM_011519808 | AGGAGCATAAGAAGGTGTCCTG | CAAGGATGTGGCATGAGCCT |
| Total α | TGGAAGCCCTGCATGCCTAC | ACTTGCTCTGAGTGGACGCT |
| β1.2 | CAGTGGGTCCTGTGATGAGG | GGACAGAGCAAGACTTCGTG |
| β1.3 | TTAAAGGAGAATGGCCCTACC | GGTATCCAGGGAACTCGTGGT |
| β1 | CCCAAAGTCACGCCCTGG | CTTCAGGAGGGCGATCTGG |
| β2 | CCCCTTCTTCTTCACCCACT | CTTCAGAAAGGACGCCCC |
| β3 | CCCAAAGTCACGGAGATCGT | CGATAGTGGATGCCTTCAGGA |
| β4 | AGGAGTCACAGTCACAGTCG | CGGCCCAGCGTGACTTTG |
| Total β | CGCTACAACCACGAGACAGAGT | GCGAGAACTCGAAGATGGGGTT |
| HPRT | AGGCGAACCTCTCGGCTTTC | TCACTAATCACGACGCCAGGG |

**Table S5. LXRα peptides detected by S-trap column coupled with MS in MDA.MB.468 cell line control samples.**

Amino acids position number based on LXRα1. Amino acid numbering position with "-" indicates the additional amino acid(s) coming before α1 and/or β1's amino acid position number 1.

| Sample | Total identified peptides | -10lgP [LXRα] | Coverage LXRα peptides | Supporting peptides [LXRα] | Unique | Amino acids position start | Amino acids position end |
|---|---|---|---|---|---|---|---|
| MDA.MB.468 siCON | 4512 | 42.09 | 12% | KC (+57.02)RQAGMR.E | No | 158 | 164 |
| | | | | R.EEC(+57.02)VLSEEQIR.L | No | 165 | 175 |
| | | | | W.PMAPDPHSREAR.Q | No | 239 | 250 |
| | | | | K.TSAIEVMLLETSR.R | No | 292 | 304 |
| | | | | YN (+.98)PGSESITFLK.D | No | 307 | 317 |
| MDA.MB.468 siLXRα | 3726 | 37.87 | 8% | R.ASSPPQILPQLSPEQ(+.98)LGMIEK.L | No | 196 | 216 |
| | | | | G.MIEKLVAAQ(+.98)Q(+.98)QC(+57.02)NR.R | No | 213 | 226 |
| | | | | R.AMNELQLN(+.98)DAEFA.L | No | 345 | 357 |
| | | | | R.M(+15.99)LMKLVSLR.T | No | 407 | 415 |
| MDA.MB.468 siLXRβ | 3641 | 44.90 | 12% | A.QGGSSC(+57.02)ILR.E | No | 33 | 41 |
| | | | | H.SAGGTAGVGLEAA.E | No | 49 | 61 |
| | | | | L.LTRAEPPSEPTEIRPQ(+.98)K.R | No | 67 | 83 |
| | | | | R.KC(+57.02)RQ(+.98)AGMREEC(+57.02)VLSEEQ.I | No | 157 | 173 |
| | | | | R.ASSPPQILPQLSPEQ(+.98)LGMIEK.L | No | 196 | 216 |
| | | | | G.MIEKLVAAQ(+.98)Q(+.98)QC(+57.02)NR.R | No | 213 | 226 |
| | | | | K.LVAAQQQ(+.98)C(+57.02)N(+.98)R.R | No | 217 | 226 |
| | | | | K.AGLQ(+.98)VEFINPIFE.F | No | 329 | 341 |
| | | | | K.AGLQVEFINPIFEFSR.A | No | 329 | 344 |
| | | | | Q.VEFINPIFEFSR.A | No | 333 | 344 |
| | | | | R.M(+15.99)LMKLVSLR.T | No | 407 | 415 |
| MDA.MB.468 treated with GW3965 | 4013 | 58.14 | 19% | R.ASSPPQILPQLSPEQ(+.98)LGMIEK.L | No | 196 | 216 |
| | | | | R.AMNELQLN(+.98)DAEFA.L | No | 345 | 357 |
| | | | | R.EDQ(+.98)IALLK.T | No | 284 | 291 |
| | | | | R.EDQ(+.98)IALLKTSAIE.V | No | 284 | 296 |
| | | | | R.VTPWPMAPDPHSREARQQ(+.98)R.F | No | 235 | 253 |
| | | | | K.LVAAQQQC(+57.02)NR.R | No | 217 | 226 |
| | | | | F.TELAIVS.V | No | 258 | 264 |

**Table S6. LXRβ peptides detected by S-trap column coupled with MS in MDA.MB.468 cell line control samples.**

Amino acids position numbers based on LXRβ1. The grey highlight indicated unique peptides of LXR variant detected by MS.

| Sample | Total identified peptides | -10lgP [LXRβ] | Coverage LXRβ peptides | Supporting peptides [LXRβ] | Unique | Amino acids position start | Amino acids position end |
|---|---|---|---|---|---|---|---|
| MDA.MB.468 siCON | 4512 | 44.58 | 17% | R.RSVVRGGAR.R | No | 113 | 121 |
| | | | | R.YAC(+57.02)RGGGTC(+57.02)QMDAFM(+15.99)RR.K | No | 123 | 139 |
| | | | | S.EAGSQGSGEGEGVQ(+.98)LTAAQEL.M | No | 205 | 225 |
| | | | | A.QELMIQ(+.98)Q(+.98)LVAAQLQC(+57.02)NKR.S | No | 223 | 240 |
| | | | | L.QVEFIN(+.98)PIFEFSRAM(+15.99)R.R | No | 345 | 360 |
| MDA.MB.468 siLXRα | 3726 | 50.87 | 18% | G.NGPPQPGAPSSSPTVK.E | No | 16 | 31 |
| | | | | R.RSVVRGGAR.R | No | 113 | 121 |
| | | | | K.RSFSDQPKVTPWPLGADPQ(+.98)SR.D | No | 240 | 260 |
| | | | | K.QVPGFLQLGREDQ(+.98) IA | No | 287 | 300 |
| | | | | A.KQ(+.98)VPGFLQLGREDQIALLK.A | No | 286 | 304 |
| | | | | A.LQQ(+.98)PYVEALLS.Y | No | 394 | 404 |
| | | | | R.M(+15.99)LMKLVSLR.T | No | 420 | 428 |
| MDA.MB.468 siLXRβ | 3641 | - | 0% | - | - | - | - |
| MDA.MB.468 treated with GW3965 | 4013 | 58.60 | 15% | G.EGVQLTAAQELMIQ.Q | No | 215 | 228 |
| | | | | V.QLTAAQ(+.98)ELMIQ(+.98).Q | No | 218 | 228 |
| | | | | G.EGVQ(+.98)LTAAQELMIQ(+.98)Q(+.98)LVAAQLQ(+.98)C(+57.02)NK.R | No | 215 | 239 |
| | | | | Q.Q(+.98)LVAAQLQ(+.98)C(+57.02)N(+.98)KR.S | No | 229 | 240 |
| | | | | R.QQRFAHFTELAIISVQ(+.98)E.I | Yes [β1] | 264 | 280 |
| | | | | K.QVPGFLQ(+.98)LGR.E | No | 287 | 296 |
| | | | | R.EDQ(+.98)IALLK.A | No | 297 | 304 |
| | | | | K.RPQDQ(+.98)LR.F | No | 410 | 416 |
